# Robustness in regulatory networks of Epithelial Mesenchymal Plasticity as a function of positive and negative feedback loops

**DOI:** 10.1101/2021.10.08.463488

**Authors:** Anish Hebbar, Ankush Moger, Kishore Hari, Mohit Kumar Jolly

## Abstract

Epithelial-Mesenchymal plasticity (EMP) is a key arm of cancer metastasis and is observed across many contexts. Cells undergoing EMP can reversibly switch between three classes of phenotypes: Epithelial (E), Mesenchymal (M), and Hybrid E/M. While a large number of multistable regulatory networks have been identified to be driving EMP in various contexts, the exact mechanisms and design principles that enable robustness in driving EMP across contexts are not yet fully understood. Here we investigated dynamic and structural robustness in EMP networks with regards to phenotypic distribution and plasticity. We use two different approaches to simulate these networks: a computationally inexpensive, parameter-independent continuous state space boolean model, and an ODE-based parameter-agnostic framework (RACIPE), both of which yield similar phenotypic distributions. Using perturbations to network topology and by varying network parameters, we show that multistable EMP networks are structurally and dynamically more robust as compared to their randomized counterparts, thereby highlighting their topological hallmarks. These features of robustness are governed by a balance of positive and negative feedback loops embedded in these networks. Using a combination of the number of negative and positive feedback loops weighted by their lengths and sign, we identified a metric that can explain the structural and dynamical robustness of these networks. This metric enabled us to compare networks across multiple sizes, and the network principles thus obtained can be used to identify fragilities in large networks without simulating their dynamics. Our analysis highlights a network topology-based approach to quantify robustness in multistable EMP networks.

**Significance Statement:** Epithelial-Mesenchymal plasticity (EMP) is a key arm of cancer metastasis. Despite extensive intra- and inter-tumor heterogeneity, the characteristics of EMP have been observed to be robust across multiple contexts. We hypothesize that topology of EMP regulatory networks contributes towards this robustness. Here, we measure the robustness of EMP in the form of its phenotypic heterogeneity and multistability and show that EMP networks are more robust to dynamical (change in kinetic parameters) and structural (change in network topology) perturbations as compared to their random network counterparts. Furthermore, we propose a network topology-based metric using the nature and length of feedback loops that explains the observed robustness. Our metric hence serves to quantify robustness in multistable EMP networks without simulating their dynamics.

## Introduction

Robustness is an inherent property of many biological systems and a fundamental feature of evolvability [1, 2]. Robust systems can maintain their functions or traits despite a dynamically changing environment, and thus possess enhanced fitness[3]. Trait robustness is pervasive in biology throughout at many organizational levels including protein folding, gene expression, metabolic flux, physiological homeostasis, development, organism survival, species persistence, and ecological resilience [4,5,6]. At the cellular level, only a limited number of specific internal/environmental changes lead to a change in the otherwise robust cell-fate. Waddington [7] postulated that mechanisms have evolved that stabilize a phenotype against both genetic and environmental perturbation. He called this process “canalization” and contrasted it with developmental “flexibility” in which a phenotype changes, adaptively, with a change in environment. Thus, understanding the mechanisms underlying robustness is of fundamental importance.

The study of robustness is also an integral part of various engineering and socio-ecological systems.[8, 9]. Designing robust machines involves integrating specific interactions (feedbacks for example) between the individual components, forming intricate networks. Similarly, biological systems such as cells or organisms also have underlying networks, with interactions that have evolved over millions of years that can impart emergent properties such as robustness to the biological system. Examples of these networks include metabolic networks, protein interaction networks, gene regulatory networks (GRNs), etc [10, 11, 12, 13]. The complexity of the interactions in these networks leads to emergent properties, which manifest as traits of the biological system. In such networks robustness can be studied in two ways, as classified by the nature of perturbations made to the network. Structural robustness is the study of robustness of biological traits to changes in the underlying regulatory network topology such as edge deletions, caused by strong perturbations such as genetic mutations [14]. Dynamical robustness, on the other hand, is the study of robustness of traits to changes in the parameters governing the dynamics of the components or nodes of the regulatory networks (production and degradation rates, link strengths etc.) [15].

Depending upon the system of study and the question of interest, the properties for which robustness is studied might differ. Two such properties of interest in multistable cell-fate decision making networks are phenotypic distribution and plasticity. Phenotypic distribution refers to the collection of phenotypes (gene expression patterns for a transcriptional gene regulatory network, for example) that the network can exhibit under different conditions or parameters. An understanding of the factors controlling these distributions for a biological system can lead to the power of manipulating the fate of said biological system despite the ever-present heterogeneity in the system. Plasticity on the other hand, refers to the ability of the network to adapt to a changing environment by modifying its phenotype, a crucial ability in many systems such as cellular differentiation and cancer metastasis. The concepts of phenotypic plasticity and heterogeneity (distribution) are well represented by Waddington’s epigenetic landscape [7] in which valleys denote gene expression patterns (phenotype) that can switch back and forth under the influence of noise or other external perturbations **(Fig 1a)**.

**Fig. 1.**
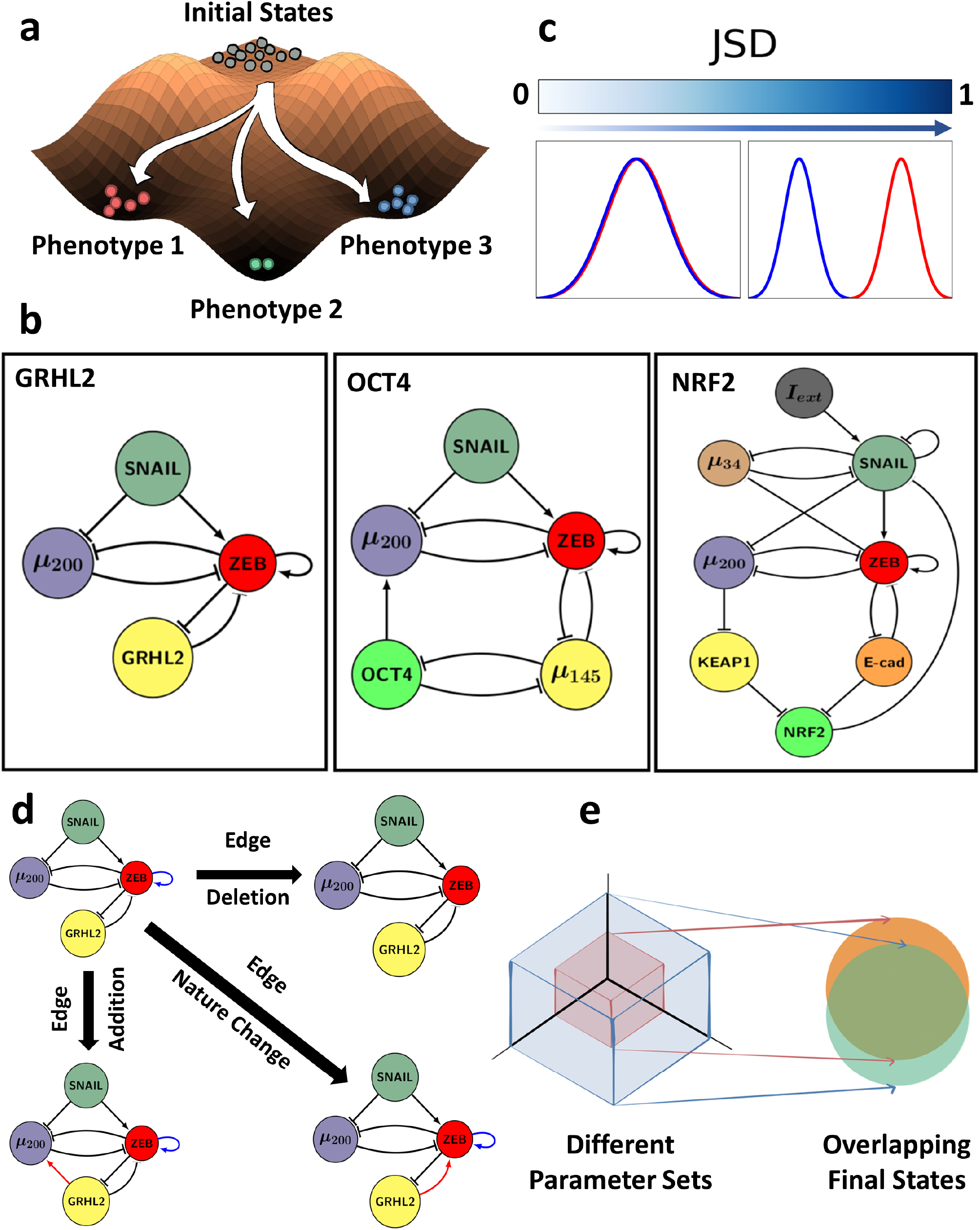
Measurement of robustness in multistable biological networks. **(a)** Depiction of plasticity as the ability of a system (defined by a set of parameters) to achieve multiple phenotypes depending on the initial state. **(b)** EMP networks analysed, namely GRHL2 (4 nodes, 7 edges), OCT4 (5 nodes, 10 edges), NRF2 (8 nodes, 16 edges). **(c)** Demonstration of how JSD between two frequency distributions varies; JSD ranges from 0 to 1. **(d)** Different ways to perturb the topology/structure of a network: deletion of an edge, addition of an edge, edge nature change from activation to inhibition and vice-versa. **(e)** Depiction of dynamic robustness where different parameter spaces converge to largely overlapping solution space.

To understand robustness, a variety of mathematical models have been used. For example, building on the concept of canalization by Waddington, Kauffman introduced a version of this concept to Boolean network modeling of gene regulatory networks by studying canalizing functions [16]. In the past decade, the mathematical frameworks for characterizing the robustness of networks have been studied in various settings, such as directed networks, embedded networks and many more [11, 17, 18]. Robustness has been widely investigated in linear control systems [19, 20] but it remains very difficult to measure the robustness of complex non-linear systems [21]. More recently, investigations of how some network topological metrics may associate with the robust behaviour of biological systems has been carried out [22, 23]. However such investigations are limited in number as well as in the scope of their generality. Hence, a comprehensive and scalable understanding of the role of network topology in robustness is required.

Here, we demonstrate the structural and dynamic robustness of the gene regulatory networks (GRNs) underlying Epithelial Mesenchymal Plasticity (EMP) **(Fig 1b)**. EMP is a fundamental cellular process implicated in development, wound healing and cancer metastasis [24, 25]. Its robust and tightly controlled dynamics have been recently investigated using quantitative experimental data and associated mathematical models [26, 27, 28]. What remains unclear though, is what factors enable the robustness of EMP. We hypothesize that this robustness is an emergent property of underlying regulatory networks. To test this hypothesis, we used a parameter agnostic, ODE based formalism: RACIPE, and a parameter independent continuous framework. We investigated the characteristics of robustness and identify underlying unifying trends. Particularly, we find that the number of positive and negative feedback loops in the system govern the extent of robustness in phenotypic distribution and plasticity. We also use this framework to predict which links in a network can be targeted and modified to inhibit plasticity – which may provide therapeutic insight in the clinic for targeting EMP. Our results thus provide an integrated understanding of the design principles that enable robustness in multistable biological networks.

## Results

### EMP networks are dynamically robust

Here, we considered various GRNs underlying EMP **(Fig 1b, S1a)**[28]. Each network has a set of activating (pointed arrows) and inhibiting (hammer-head arrows) edges/links connecting the nodes of the network. These activating and inhibiting interactions are modeled using ODE formalism, where varying strength of these interactions alter the production rates of corresponding target nodes. Here, we focus on two emergent dynamical outputs of a given GRN: phenotypic heterogeneity (frequency distribution of different discretized steady states/phenotypes) and plasticity (fraction of randomly chosen parameter sets in ODE simulations that can give rise to multiple steady states). To identify how robust these two outputs are to the underlying network topology, we perturb them in multiple ways and re-calculate the outputs of perturbed networks. The perturbations discussed here include edge additions, deletions and changing the sign of an edge from an activatory to inhibitory and vice-versa (**Fig 1d**).

The smaller the deviation in terms of these outputs of perturbed (hypothetical) networks when compared with those of wild-type (EMP) network, the higher the robustness of corresponding EMP network. We measure changes in plasticity through fold-change, and those in phenotypic distributions by using an information theoretic metric called Jensen Shannon Divergence (JSD) [29] (See Methods and **Fig 1c**). Both fold-change and JSD are defined on a scale of 0 to 1. The closer the fold-change is to 1, the higher the similarity between plasticity of the perturbed network and that of the corresponding EMP network. Similarly, the lower the JSD, the smaller the dissimilarity between phenotypic distributions emergent from the two networks: perturbed and corresponding EMP one. If a network displays similar emergent behaviour even upon perturbation of the network topology, it is said to be structurally robust. Similarly, if the behaviour of the network roughly remains the same under variation of the kinetic parameters governing each interaction, such as the production and degradation rates, regulatory interaction parameters between transcription factors and promoters, etc., it is said to be dynamically robust **(Fig 1e)**.

We simulate each network using a ODE-based framework called RACIPE [30] (see Methods). For a given GRN, the framework first constructs a set of coupled ordinary differential equations. Then, multiple sets of ODE parameters are generated from pre-determined ranges (the parameter space), which are then used to simulate interactions between nodes of the network. These simulations can be used to identify the robust dynamical features of a particular network. Because RACIPE randomly samples parameters for each simulation, it can be thought of as a parameter agnostic framework [30]. Using this framework, we can analyze the dynamic robustness of a network by studying the variation in network behavior with changes in RACIPE parameter space (dynamic perturbation). Plasticity score of the network is then defined as the fraction of such random parameter sets that can give rise to multiple stable states[28]. The phenotypic distribution is found using the frequency distribution of the discretized steady states of the network (see Methods for discretization procedure) over the RACIPE parameter space.

Because the network behavior in RACIPE is regulated by both network topology and various kinetic parameters, plasticity and phenotypic distribution can vary across distinct parameter ensembles. In order to characterize this variation with across the parameter space, we simulated these EMP networks using RACIPE, by varying the maximum ranges for different network parameters via multiplying them by a factor ranging from 1/3 to 3. While multiplication factor > 1 leads to the expansion of the parameter space, multiplication factor < 1 shrinks the parameter space. The Hill coefficient in these systems contributes to the inherent non-linearity and hence multistability [31]. We varied the default RACIPE parameters in 3 different ways: all parameters (Hill coefficient, production rate, degradation rate and fold change) being varied, everything but Hill coefficient, and only the Hill coefficient. We then calculated the JSD of the resultant phenotypic distribution from that of the unmodified parameter space and plotted them against the multiplication factors **(Fig 2a, S2a,b)** for all EMP networks. As the multiplication factor got farther from 1, the JSD increased. Visually speaking, the proportional increase was higher when the parameter space shrunk, as opposed to when the parameter space was expanded. However, the JSD values in all cases was relatively low (maximum value across networks = 0.25, see **Fig2a, inset**).

**Fig. 2.**
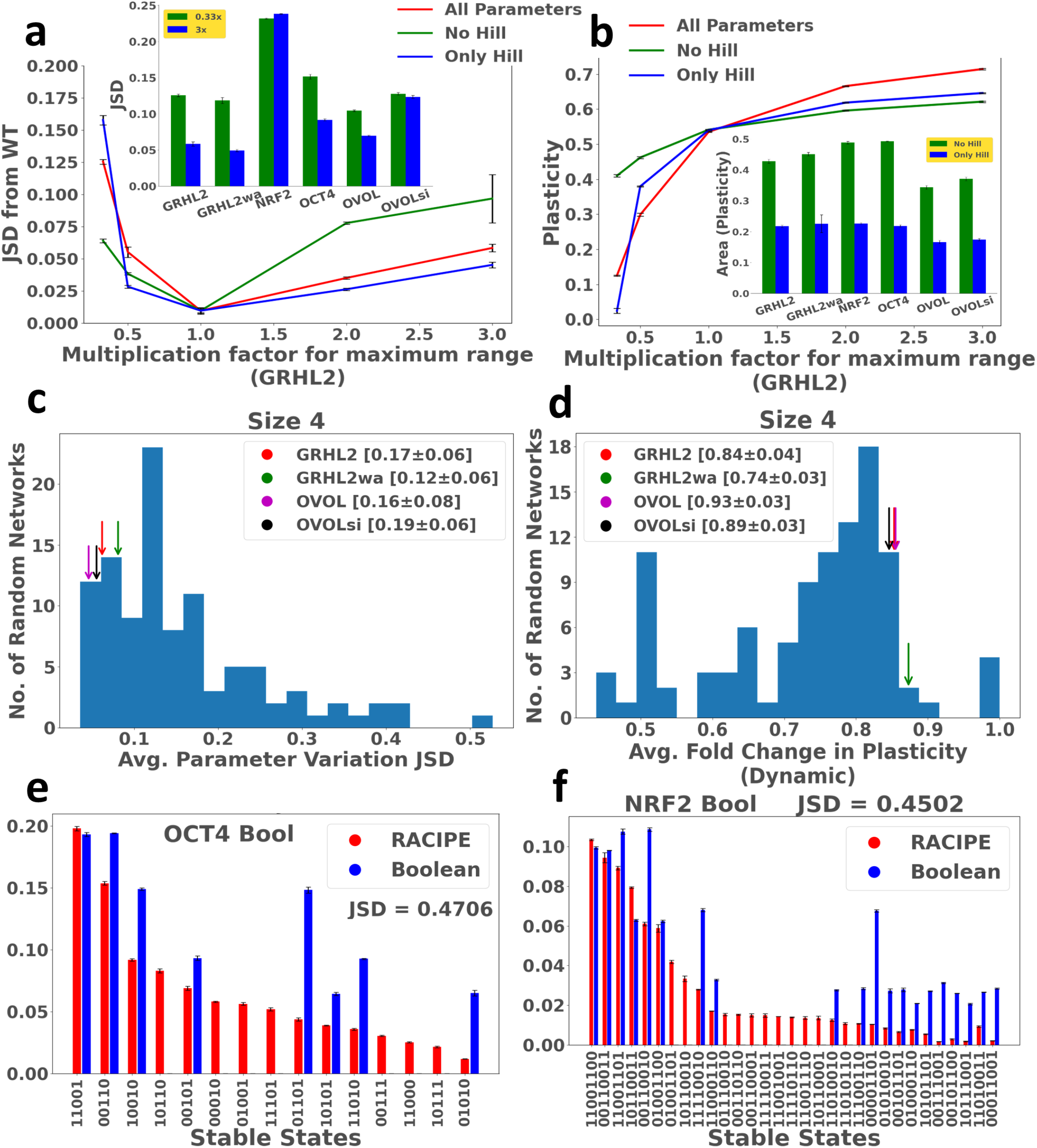
High dynamical robustness of EMP networks. **(a)** Variation of phenotypic distribution (quantified by JSD) upon scaling by the multiplication factor (x axis), the range(s) of: all parameters (red), only Hill coefficient (blue) and all parameters but Hill coefficient (red). The barplot (inset) depicts JSD at 0.33× and 3× multiplication ranges for all WT networks. Errorbars represent mean +/− standard deviation. **(b)** Similar to **a**, but for plasticity. The barplot (inset) depicts the difference between varying only hill coefficient (blue) and every parameter except hill coefficient (green). Errorbars represent mean +/− standard deviation. **(c)** Histogram of average JSD for random networks of size 4, with corresponding EMP networks marked with colored arrows. Percentiles for WT networks reported as Mean +/− standard deviation. **(d)** Same as **c**, but for average fold change in plasticity. **(e)** RACIPE (red) vs Boolean (blue) stable state distributions (OCT4), and corresponding JSD. The *y*-axis is the frequency of the stable state. **(f)**Same as **(e)**but for NRF2 network.

Similarly, we plotted the plasticity of the perturbed parameter space against the corresponding multiplication factor **(Fig 2b, S2c,d)**. As the parameter space increased, the plasticity of the network increased as well (as we are allowing for more multistable parameter sets). The range of plasticity in this case was much larger ranging from 0.1 – 0.7 in some cases. We then compared the absolute area between the control (all parameter variation), with the only hill coefficient variation and no hill coefficient variation curves (See Methods) in order to quantify the closeness of the regimes. The dynamic fold change in plasticity was affected mainly by the variation in Hill coefficient **(Fig 2b, inset)**, suggesting that the Hill coefficient (non-linearity) is a crucial parameter in enabling multistability.

Next, we asked whether this dynamical robustness is specific to these network topologies. To answer this, we generated three sets of 100 random (hypothetical) networks corresponding to each network size of EMP networks, having the same number of nodes as the EMP networks, but randomly placing activating/inhibiting edges to connect the nodes (see Methods). In this context, we refer to the EMP networks as “wild-type” (WT) networks. We then measured the dynamical robustness of these networks in the same manner as the WT networks, i.e., using the measures of average fold change in plasticity (see Methods) and JSD upon varying multiplication factor, and the analysis was repeated in triplicates.The distribution of these values, when plotted, revealed that WT networks had much lower JSD, and an average fold change in plasticity much closer to 1 than most of the random networks **(Fig 2c,d, S2e,f)**. Moreover, the percentiles of the WT networks in the histograms were consistent across all 3 runs (statistics shown in **Fig 1c,d**), further reinforcing that the EMP networks are more dynamically robust than their random counterparts.

### A continuous, parameter independent formalism for efficient simulation of GRNs

Another way to understand the variation due to kinetic parameters is to compare the network behavior in RACIPE simulations to that of a parameter independent framework, such as the Boolean (logical) formalism [32]. The dynamics of Boolean models are governed purely by the topology of the network. The gene activities are discrete (either ON or OFF), which gives us a more qualitative understanding of how the nodes in the network influence each other to give rise to varying network dynamics. Therefore, the Boolean formalism can serve as a non-parametric counterpart to the RACIPE formalism. One drawback however is that with the Boolean formalism, we cannot define plasticity the way we define it for RACIPE, therefore, only robustness in phenotypic distribution can be studied.

We applied a threshold-based Boolean formalism [32] to simulate these networks (see Methods). While the top states obtained from both formalisms are the same, dissimilarity between the steady state distributions obtained in both frameworks is relatively large (*JSD* ~ 0.5 **Fig 2e,f, S2g,h**). While this range of JSD values could indicate lack of dynamic robustness in EMP networks, we asked if the high dissimilarity between RACIPE and Boolean might be caused by fundamental differences between the methods, other than the lack of parameter sets in the latter.

The existing Boolean formalism differs from RACIPE in two aspects: the lack of kinetic parameters as compared to RACIPE and a discrete state space instead of a continuous one. Thus we decided to modify the Boolean formalism by making its state space continuous **(Fig 3a)**. The details of the implementation of this formalism, termed Continuous State-Space Boolean (CSB) formalism, are given in the Methods section. This formalism provides a better measure of the dynamic robustness of the networks, because the essential difference between the CSB formalism and RACIPE that is now left is the lack of kinetic parameters in the former, so any differences that arise can be thought of as coming from these kinetic parameters. While this formalism is parameter independent, the steady state phenotypic distributions obtained in this model are much closer to that obtained via RACIPE (lower JSD), relative to the earlier Boolean model, for all EMP networks **(Fig 3b, S3b,c)**. Most random (hypothetical) networks showed this increased similarity as well **(Fig S3a)** suggesting that this framework can be applied to a larger set of networks. Moreover, this model is computationally inexpensive compared to RACIPE as measured for simulations across varying network sizes **(Fig 3c)**. We also investigated the sensitivity of the results of both frameworks to the number of simulations, namely analysing how much the output phenotypic distribution varies across different simulations for the same number of total simulations (see Methods). We see that the results obtained in the CSB are more consistent for the same number of simulations when compared to RACIPE, further highlighting its benefits as an alternative to ODE based methods **(Fig 3d)**.

**Fig. 3.**
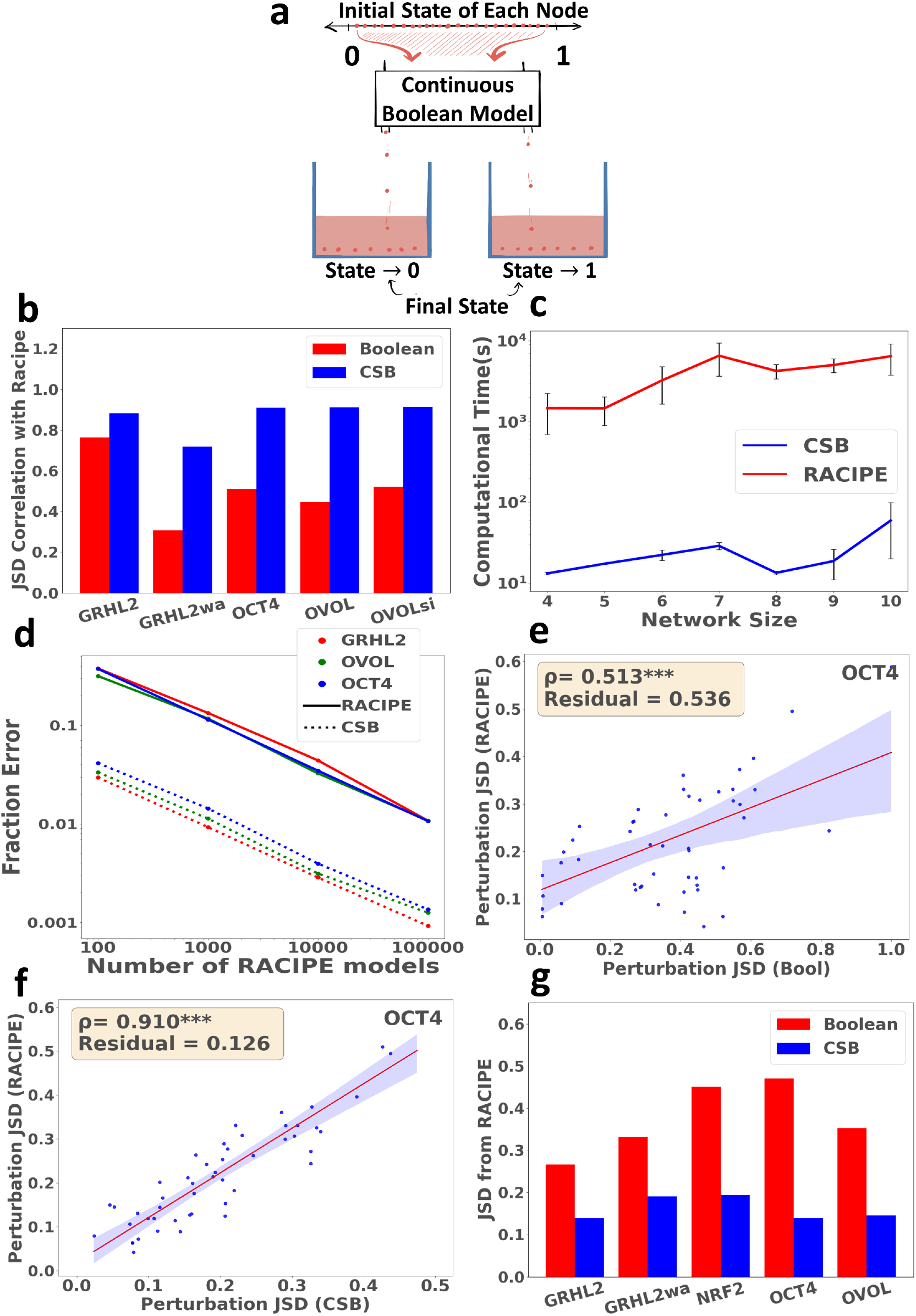
Continuous State Space Boolean (CSB) formalism performs better than the Boolean formalism. **(a)** Visualization of the CSB framework. Networks are simulated in a continuous state space and the resultant final states are discretized. **(b)** Comparison of JSD of the distributions obtained by RACIPE from Boolean framework (red) and CSB framework (blue) for various EMP networks. **(c)** Comparison of computation times for RACIPE and CSB for networks of different sizes. Error bars represent mean +/− STD of computation time, obtained by simulating 5 random networks of each size. **(d)** Comparison of error in phenotypic frequency for RACIPE and CSB simulations of WT networks. **(e)** Comparison of perturbation JSD obtained via RACIPE (y-axis) and Boolean (x-axis) for OCT4 network. Each dot represents one perturbed network. **(f)** Same as **e**, but with CSB perturbation JSD on x-axis. **(g)** Grouped barplot comparing the correlation of JSD of RACIPE perturbations from WT vs JSD of Boolean/CSB perturbations from WT for various EMP networks. Statistical significance for correlations is reported : *** - *pvalue* < 0.001

We then sought to understand if the JSD due to structural perturbations, i.e. either a single edge deletion, insertion, or a nature change **(Fig 1d)**, in the CSB correlated well with the same perturbation done in RACIPE. We calculated the JSD of the perturbed network’s phenotypic distribution from that of wildtype networks, using RACIPE, Boolean and CSB fomalisms. We found that the correlation between RACIPE and CSB JSD values, as well as the residual obtained via linear regression, are much better than that between RACIPE and Boolean **(Fig 3e-g, S3d-i)** across the EMP networks examined, further providing support to the similarity of the CSB to RACIPE. We then simulated random networks using the CSB formalism to check if the dynamical robustness properties are captured by this formalism. The JSD between RACIPE and the CSB formalism for the phenotypic distributions of EMP networks was lower than that of most random networks **(Fig 4a, S4a,c)**. Consequently, the CSB proved to be a framework that is both computationally efficient and correlates well with RACIPE perturbations of the corresponding network topology. This result suggests the importance of network topology in determining the phenotypic distribution. While individual parameter sets do have unique effects on the emergent phenotypes, the overall phenotypic distribution can still be predicted using the network topology alone.

**Fig. 4.**
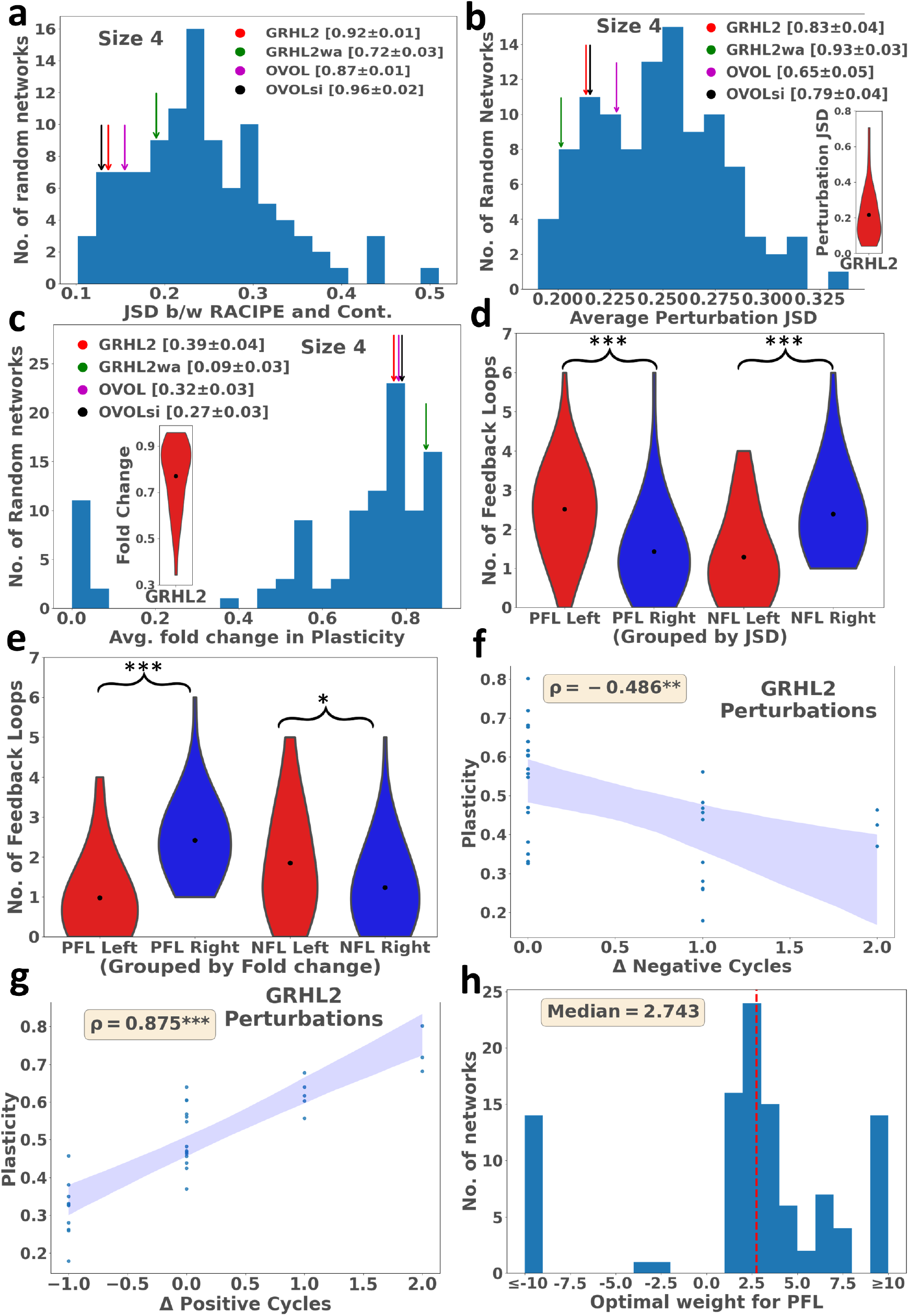
Structural robustness and feedback loops. **(a)** Distribution of the JSD b/w the phenotypic distributions from RACIPE and CSB for random networks of size 4, with the corresponding WT networks marked. Percentiles for WT networks reported as Mean +/− standard deviation. **(b)** Distribution of the average perturbation JSD of random networks of size 4, with the corresponding WT networks marked. The violin plot (inset) indicates the distribution of the perturbation JSDs for GRHL2. Percentiles for WT networks reported as Mean +/− standard deviation. **(c)** Same as **b**, but for fold change in plasticity, **(d,e)** Violin plots depicting the distribution of negative and positive feedback loops on either side of the median of the histograms **b,c**. Unpaired T-test comparing the distribution of feedback loops on either side of the median was performed, with significance depicted on top of the plots. **(f)** Scatter plot of the plasticity of the perturbed network vs the change in PFLs for GRHL2. Each dot represents a perturbed network. **(g)** Same as **f**, but for NFLs. **(h)** Histogram of the optimal weights for PFLs for random and WT networks (Size 4). Statistical significance for correlation and t-test is reported as follows: * – *pvalue* < 0.05,** – *pvalue* < 0.01,*** – *pvalue* < 0.001.

### Structural robustness of multistable EMP networks

We further investigated the robustness of plasticity and phenotypic distribution to structural perturbations, i.e., addition, deletion and nature change of network edges. For robustness in phenotypic distribution, we simulated the random networks and their perturbations using the CSB formalism. We then calculated the JSD of each perturbed network from its corresponding unperturbed network **(Fig 4b inset)** and took an average of these JSD values to represent the structural robustness of the network. The distribution of the average perturbation JSD values for random networks revealed that the EMP networks have a lower average perturbation JSD and therefore higher structural robustness in phenotypic distribution as compared to most random networks when perturbed similarly **(Fig 4b, S4b,d)**. However, single edge perturbations may not fully capture the robustness landscape, as it is also possible to modify multiple edges at the same time. Consequently we looked at multiple random edge perturbations in the EMP networks **(S4e,f)**. If the graph had E number of edges, we performed up to E number of random edge perturbations to see how the average JSD varies with perturbation size. We observed that the average perturbation JSD for a single edge was about half of the average obtained in E many perturbations, indicating that analysing the effects due to a single perturbation is sufficient to understand the phenotypic robustness of a network to edge modifications. Hence, the CSB formalism is a useful tool to investigate the structural robustness of a network, as the average value of the perturbation JSDs such obtained are a good indicator of the global network robustness in phenotypic distribution.

For the analysis of structural robustness in plasticity, we simulated the random networks and the corresponding perturbations using RACIPE, because plasticity is defined only for RACIPE. Similar to JSD, we calculated the average fold change in plasticity for each random network and obtained a distribution of the same. We found that the average fold change in plasticity, obtained from a distribution of the fold change in plasticity upon all topological perturbations (**Fig 4c, inset**) of WT (Wild Type) networks is closer to 1 than that of most random networks **(Fig 4c, S4g)**, indicating that the EMP networks are structurally more robust than their random network counterparts.

### Feedback loops underlie the robustness in EMP networks

After characterizing structural robustness in the EMP networks, we attempted to understand why these EMP networks show structural robustness. Positive feedback loops have been reported to play a major role in the stability of biological networks by reinforcing the network such that the nodes involved in the loops are inclined to display similar expression levels [33, 34, 35]. The reinforcement provided by positive feedback loops can lead to convergence to approximately similar steady states across a wide range of parameters in RACIPE. Moreover, positive feedback loops have been shown to play a crucial role in governing plasticity of the networks as well [28]. Similarly, negative feedback loops can contribute to oscillations [33, 35, 34]. Therefore, we hypothesized that positive and negative feedback loops can govern the robustness of these GRNs.

To understand if the positive and negative feedback loops play a role in imparting structural and dynamical robustness to networks, we used the ensemble of random networks created. First, we divided the distributions of average perturbation JSD **(Fig 4b)** and average fold change in plasticity **(Fig 4c)** about their respective medians, and asked if there was a significant difference in the distribution of feedback loops for the networks with high robustness vs those with low robustness. The networks with the lower average perturbation JSD had a significantly higher number of positive feedback loops (PFLs) and a lower number of negative feedback loops (NFLs), as quantified by the difference in the means of the distributions **(Fig 4d)**. Interestingly, the shift in the distribution of negative feedback loops is more distinct when compared to that of positive feedback loops. Similarly the networks with a higher average fold change in plasticity have a smaller number of negative feedback loops and a larger number of positive feedback loops **(Fig 4e)**. Unlike with JSD, the two groups of networks in case of plasticity are better differentiated by positive feedback loops than with negative feedback loops. These results suggest that positive and negative feedback loops might have varying degree of importance in governing different kinds of robustness. A common feature for robustness in both plasticity and JSD seems to be, however, that the networks with higher robustness have a larger number of positive feedback loops and a smaller number of negative feedback loops.

To better understand the relative contribution of positive and negative feedback loops, we went back to EMP networks and measured the Spearmann correlation between plasticity of the perturbed networks and the corresponding change in number of positive and negative feedback loops (**Fig 4f,g**, each dot is a perturbed network). As expected, the number of positive feedback loops has a stronger (~ 2 fold in magnitude) correlation with the fold change in plasticity in comparison to the number of negative feedback loops. This observation suggests that positive and negative feedback loops have opposite but unequal effect on network plasticity. We then investigated if a weighted combination of positive and negative feedback loops can help explain plasticity better. For a given network, we identified the optimal weights for positive and negative feedback loops that would maximize the correlation between the combined loops metric and plasticity. The larger the value of the weight, the higher is the contribution of the corresponding loops in explaining plasticity (see methods section). Because PFLs correlate positively with robustness in plasticity and NFLs correlate negatively, we assigned a negative weight to NFLs (robustness should decrease as their number increases) and a positive weight to PFLs (robustness should increase as their number increases). The weight distribution obtained from the ensemble of random networks had a median of around 3 : −1 **(Fig 4h)**, i.e, positive cycles were given more weight in explaining plasticity. As expected, we find that the sign-weighted feedback loops (SWFL = weighted sum of PFLs and NFLs, see Methods) has a better correlation with the plasticity of perturbed networks, as compared to just the PFLs **(Fig S5a)**.

Previous literature also suggests that not all PFLs play the same role in regulating the plasticity of the network[36, 37]. Consequently, we decided to adopt another independent weighting strategy: penalize each PFL using its length, i.e., giving a higher importance to shorter feedback loops (see Methods), with the hypothesis that breaking a longer feedback loop would have a more diluted effect on the system. We then took a sum over these penalized PFLs, creating a new metric (WPFL). To verify that this penalization is an improvement on the results obtained without penalization, we compared the correlations obtained via the two metrics (Δ*PFL* and Δ*WPFL*) with the plasticity of the network obtained via structural perturbation for the ensemble of random networks **(Fig 5a)**. The correlation using WPFLs is an improvement on PFLs (most dots are above the *x* = *y* line), indicating that it is important to consider the length of the feedback loops as a factor influencing their impact.

**Fig. 5.**
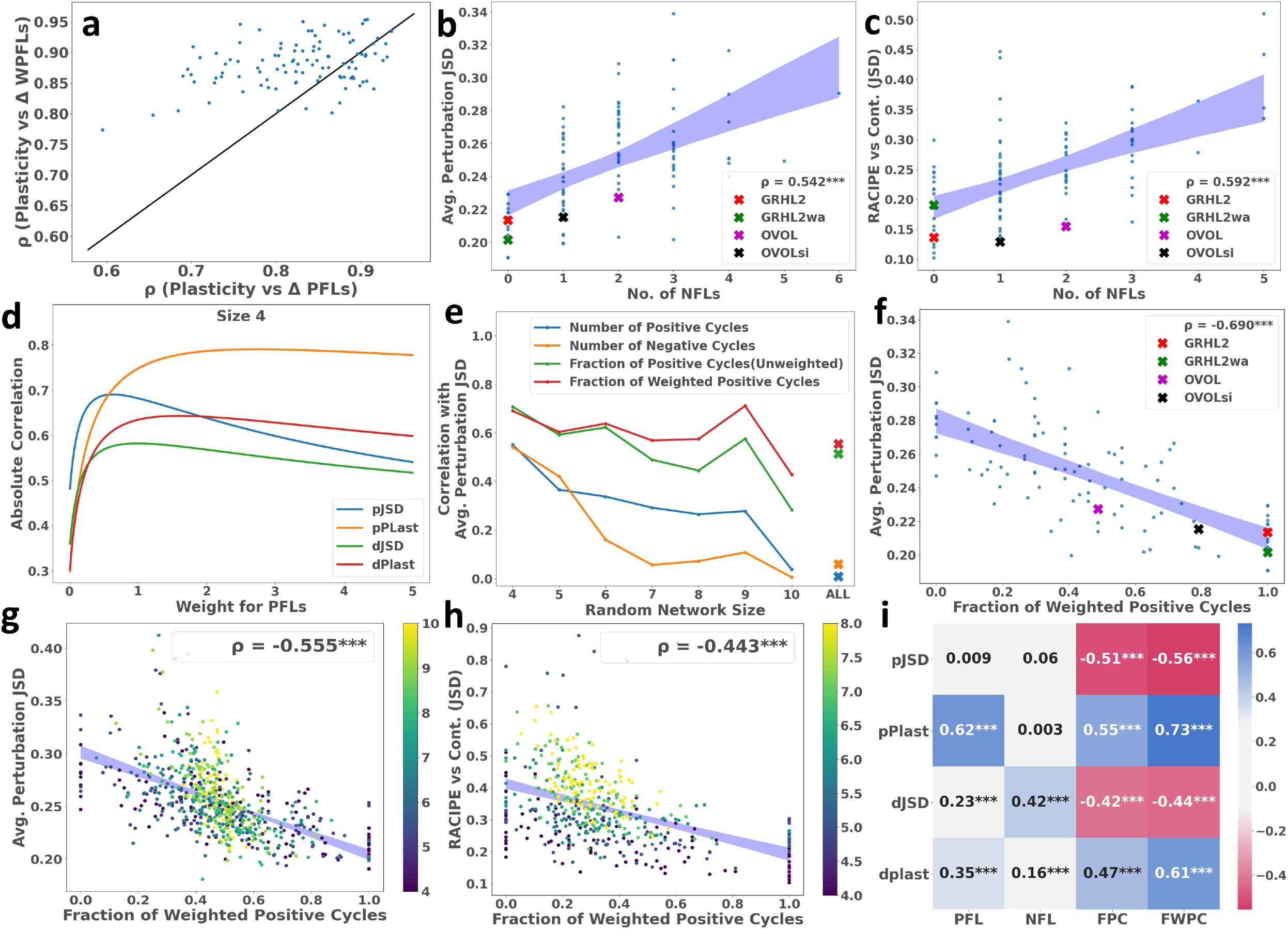
Fraction of weighted positive feedback loops correlates with network robustness. **(a)** Scatter plot of the correlation of fold change in plasticity with positive feedback loops and weighted PFLs, along with the *x* = *y* line. **(b)** Scatter plot of the average perturbation JSDs from WT vs No. of negative feedback loops for networks of size 4, with the corresponding WT networks marked. **(c)** Same as **b**, but for JSD b/w Racipe and CSB. **(d)** Change in the strength of correlation between the robustness measures-structural robustness in JSD (pJSD), plasticity (pPlast), dynamic robustness in JSD (dJSD), and plasticity (dPlast)- and fraction of weighted positive feedback loops (FWPC), with change in the weight of WPFL in FWPC, for networks of size 4. **(e)** Correlation b/w Avg. perturbation JSD and PFLs, NFLs, fraction of positive cycles (FPC), and FWPC for networks of different sizes. Absolute values have been used for ease of comparison. **(f)** Average perturbation JSD from WT vs the fraction of weighted positive cycles for networks of size 4, with the corresponding WT networks marked. **(g)** Scatter plot of the Average perturbation JSD from WT vs the fraction of weighted positive cycles for networks of sizes 4-10, coloured by size. **(h)** Similar plot as **f** for the JSD b/w Racipe and Continuous stable state frequencies for random networks of sizes 4-8. **(i)**Heatmap of the correlation coefficients of robustness measures (pJSD, pPlast, dJSD, dPlast) with PFLs, NFLs, FPC and FWPC. Statistical significance of correlation is reported as follows: *** –*pvalue* < 0.001

We then decided to check if feedback loops correlate with the other aspects of robustness as well. We found that the average perturbation JSD correlates positively with the NFLs **(Fig 5b)** and negatively with PFLs (**Fig S5b**) for all random networks of size 4, and the JSD between RACIPE and Continuous also correlates positively with NFLs **(Fig 5c)**. These trends reiterate that structural robustness correlates negatively with the NFLs and positively with PFLs. These trends are characteristic of the sign of the feedback loops, as the correlation of structural robustness with total number of feedback loops (PFL+NFL) was weak and not significant (**Fig S5c**).

However, PFLs make for a poor metric for predicting the robustness of a network. As there are no clear bounds on the number and no direct dependence on the network size, the correlation between PFLs and robustness does not hold when networks of multiple sizes are compared **(Fig S5d)**. One alternative is hence to consider the fraction of PFLs 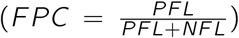 instead, as it is bounded between 0 and 1 irrespective of the network size. From abovementioned results, we also know that both the length and sign of the feedback loop are important factors to be considered. Thus, we decided to look at both weighting strategies simultaneously: the fraction of weighted positive feedback loops (FWPC) instead, where positive and negative feedback loops are first weighted by length, and then weighted by sign so as to maximize the correlation with robustness (see Methods). Note, that these weights are calculated separately for each measure of robustness (**Fig 5d**). While the optimal weights are taken as the ones that maximize the absolute value of the correlation, in most cases, the decrease in the correlation for weights greater than the optimum weight is negligible (**Fig S6a-d**). The optimum weights for each robustness measure-network size combination is given in **Table S1**. Upon comparing the correlation coefficients for average Pertubation JSD vs the different metrics mentioned above (PFLs, NFLs, FPC, FWPC) for different network sizes (4-10, **Fig 5e**), we find that the correlation against PFLs and NFLs drops as the network sizes increase, especially when networks of different sizes are considered simultaneously (**Fig 5e**, labelled “ALL” on x-axis). On the other hand, the fraction of weighted positive cycles consistently has a higher correlation as compared to the other metrics. On plotting the average perturbation JSD for random networks of various sizes vs FWPC **(Fig 5f,g)**, we see that the correlation is maintained even across multiple network sizes, giving us a scale-independent understanding of the structural robustness of networks.

FWPC also correlated positively with structural and dynamic robustness in plasticity, and dynamic robustness in distribution as measured by JSD between Continuous and RACIPE **(Fig 5h, S5e,f)**. Furthermore, we found that the random networks for which the JSD between RACIPE and the CSB was higher compared to Boolean were characterised by having a smaller fraction of positive feedback loops (FPC) (**Fig S5g**), supporting the connection between the fraction of positive feedback loops and degree of dynamic robustness. It is, however, interesting to note that while the biological networks we studied had a high dynamic robustness in distribution as measured by the JSD from RACIPE parameter variation as well, this robustness did not correlate strongly with the fraction of weighted positive cycles **(Fig S5h)**.

The correlation coefficients obtained using the metrics above for different robustness criteria, combining networks of all sizes is shown as a heatmap **(Fig 5i, S6e,f)** (done in triplicates). Note the higher correlation when using the FWPC, as opposed to the NFLs, PFLs, or the FPC in these networks. Based on these results, we can conclude that the fraction of weighted positive feedback loops serves as a good metric to measure both structural and dynamic robustness in networks. Because the calculation of this metric does not require any simulations, these metrics have extreme computational value for determining the robustness of large-scale biological networks.

While the optimally weighted FWPC metric explains robustness measures well, the drawback with the metric is its lack of generality, i.e., the optimal weight must be calculated for each robustness measure and network size combination. Such calculations lose feasibility as the network size increases. Hence, we sought to generalise the metric. A closer look at the change in absolute correlation with the weight **(Fig 5d, S6a-d)** revealed that, at the weight ratio of 1 : −1 (between WPFLs and WNFLs) the correlation between FWPC and robustness metrics is close to the maximum possible. Hence, we decided to use the weight of 1 : −1 for the calculation of FWPC for any given network as a default.

### Larger regulatory networks follow similar design principles of robustness

To test the scalability of our results, we analysed 2 large EMP networks: EMT RACIPE (22 nodes, 82 edges)[38] and EMT RACIPE2 (26 node, 100 edges)[39] **(Fig 6a,b)**. We first found the phenotypic distributions of these networks in all 3 models, RACIPE, Boolean and CSB **(Fig S7a-d)**. As it was with the small networks, the CSB showed a better agreement with RACIPE than Boolean. We also found the distribution of multistability (parameter sets giving rise to n steady states, n = 1,2,3…) in RACIPE for these networks **(Fig 6c,d)**, via 100, 000 simulations in triplicates. We observed that these networks have very high plasticity, with only upto 5% of parameter sets displaying monostability, the rest being multistable (See Methods). Note that the error bars in these simulations are very small, showing that the number of simulations performed give an accurate depiction of the fraction of multistable states. We then perturbed these networks structurally (single edge deletions/ nature changes, 144 and 202 perturbations respectively) to study their robustness in plasticity. Both of these networks have a large number of positive feedback loops (> 3000, > 10000 respectively), and most perturbations only caused minor change in the plasticity score, thereby demonstrating the structural robustness of these networks. We found the plasticity scores for each of the perturbed networks, and as observed in the networks of smaller scale, the correlation between the plasticity of the perturbed network, and the corresponding change in the number of positive feedback loops was positive and significant **(Fig 6e,f)** (See Methods for details on these metrics)

**Fig. 6.**
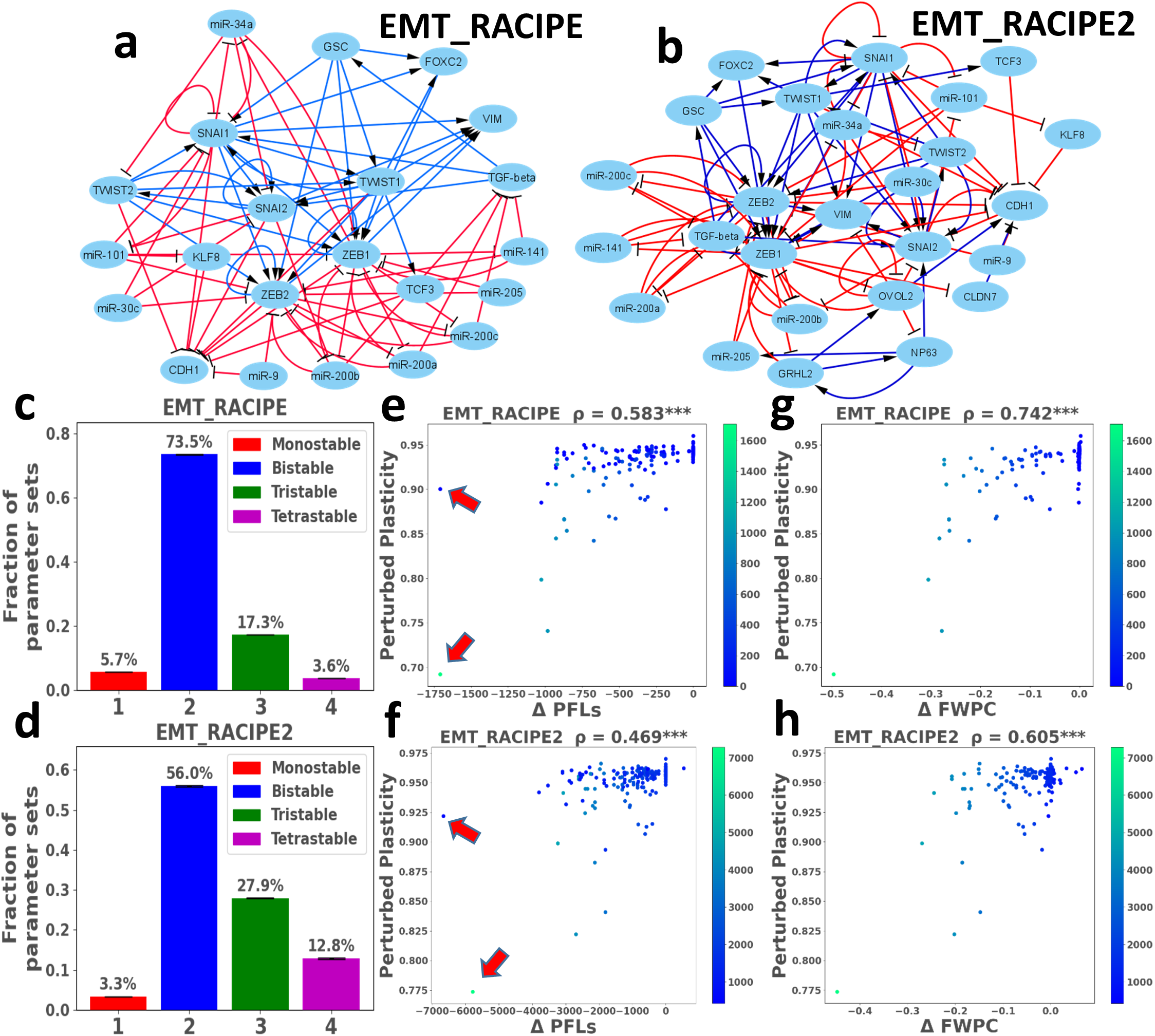
Feedback loop based metric explains robustness in large-scale networks. **(a,b)** The 22 node EMP network (EMT_RACIPE) and 26 node EMP network (EMT_RACIPE2). **(c,d)** Barplot of the fraction of multistable parameters in RACIPE for EMT_RACIPE,EMT_RACIPE2. **(e,f)** Plasticity of the perturbed network vs change in positive feedback loops for each perturbed network (EMT_RACIPE,EMT_RACIPE2) was plotted. **(g,h)** Same as **e,f**, but for change in FWPC. Color code denotes number of negative feedback loops in a given perturbed network topology. Significance of correlation is reported: *** − *pvalue* < 0.001

However, in both networks there were two perturbations (**Fig 6e,f**; marked with red arrows) that reduced the number of positive feedback loops equally drastically, yet have quite distinct fold change in plasticity. Consequently, it is clear that just the change in the number of positive feedback loops alone does not provide a complete picture of how the plasticity of a network changes upon perturbations. As our results show that negative feedback loops also matter in determining the plasticity, we also coloured each perturbation on the plot by the number of negative feedback loops. It was then seen that among the two perturbations, the one with a lower fold change in plasticity had a higher number of negative feedback loops, confirming that negative feedback loops play an important role in regulating the plasticity of a network. We then plotted the fold change in plasticity against the change in FWPC (FWPC = WPFL/(WPFL+WNFL), as positive and negative loops were given the same weight), and found that the metric showed stronger correlation with plasticity in comparison to the number of positive feedback loops **(Fig 6g,h)**. These results hence suggest that the fraction of weighted positive cycles (FWPC) can be used to understand the robustness (and fragility) of larger networks as well.

To establish a causal connection between FWPCs and structural robustness of networks, we perturbed the NRF2 network and a few random networks of same size as that of NRF2 (but with altered topology, as a control case). For each network, we identified the edge perturbations that led to the maximum increase or decrease in FWPC for these networks. We then continued to make iterative perturbations that lead to further increase in FWPC value until it reached 1 (or decrease in it until it reached 0) (**Fig 7a,b**). Our results suggest that reduction in FWPC associates with lower network robustness and vice-versa. We measured the structural robustness by calculating the JSD (using the CSB model) for each of these perturbations in both NRF2 and multiple random networks. We found a monotonous decrease in perturbation JSD (PJSD) with and increase in FWPC (**Fig 7c,d**) both for NRF2 and for a representative random network. Similar monotonic trends between structural robustness and FWPC were observed for other random networks of 8 nodes as well (**Fig 7e,f**). These results strengthen our previously shown correlation-based associations that perturbations made to increase the FWPC metric make the network more robust.

**Fig. 7.**
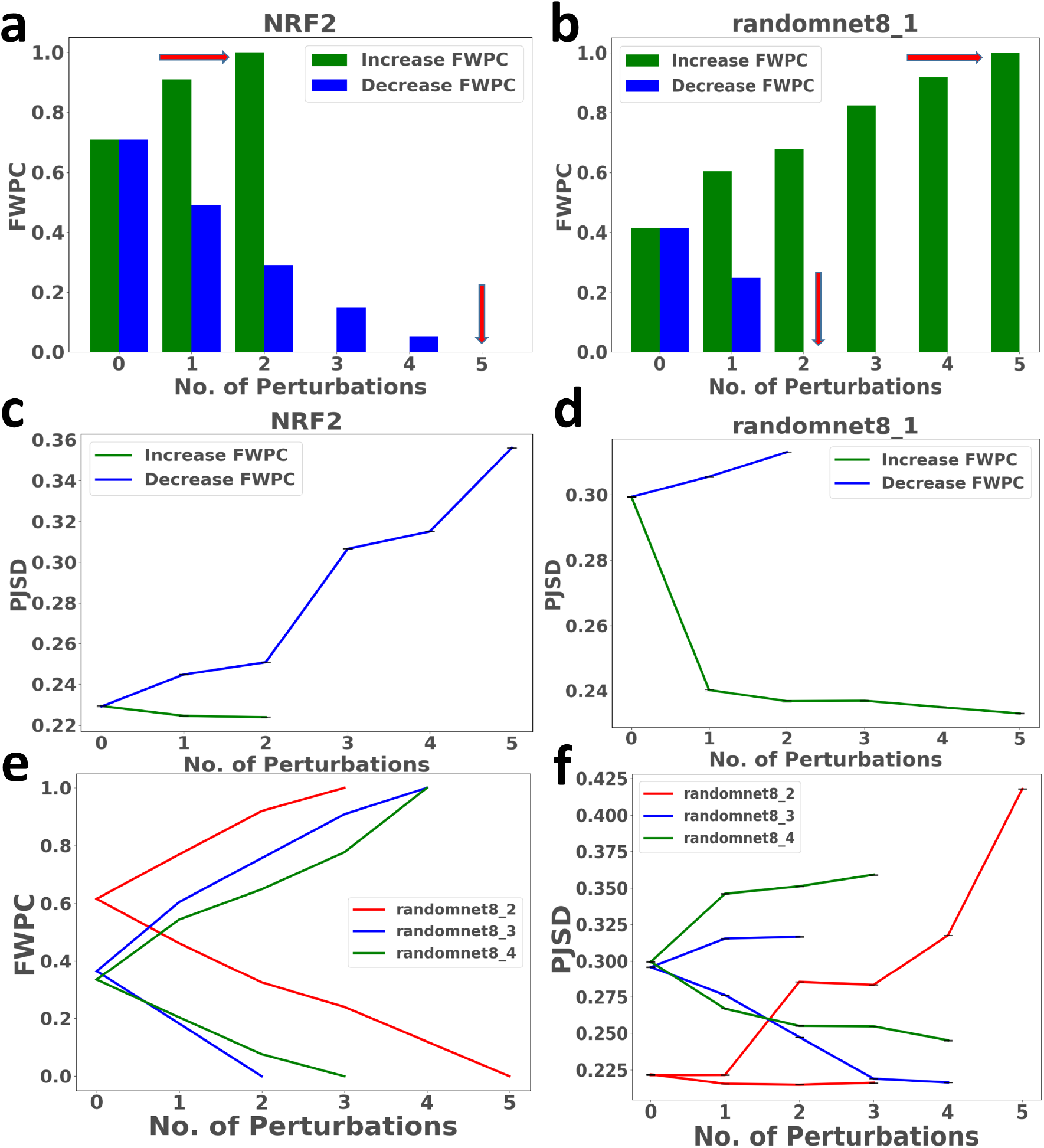
Perturbations reducing FWPC reduce network robustness and vice versa. **(a,b)** Representative barplot showing the edge perturbations that increase or decrease FWPC maximally iteratively. The red arrows show the point at which perturbations were stopped, when either the maximum (1) or minimum (0) FWPC were reached. **(c,d)** Change in perturbation JSD for NRF2 and a representative random network of size 8 with each perturbation described in panels a and b. **(e)** Change in the FWPC metric with network perturbation for three additional random networks. **(f)** Change in structural robustness (perturbation JSD) with network perturbation for three additional random networks. In panels **d, f**; mean +/− standard deviation across three replicates are reported.

## Discussion

Robustness is a fundamental, ubiquitous feature of biological systems, that enables them to maintain their function in fluctuating environments against both dynamic and structural perturbations. Decoding what network motifs can protect network dynamics against such perturbations is key to understanding robustness. Our results provide a theoretical toolset to identify robustness in both phenotypic distribution and plasticity in multistable EMP networks using a network topology based approach. We have simulated multiple EMP networks along with an ensemble of their random network counterparts in order to identify the design principles that enable these networks to display robustness when subjected to dynamic or structural perturbation. We also pave the way towards understanding how one can perturb the fragile components of these networks to cause a change in the degree of multistability (plasticity). Our analysis shows that reducing the number of positive feedback loops and/or increasing the number of negative feedback loops in networks can alter their plasticity across a wide range of parameter sets. A recent study complements our conclusion by showing that disrupting the miR-200/ZEB positive feedback loop, one of the key driving factors of EMP, via CRISPR led to a significant drop in metastasis *in vivo* [40].

EMP dynamics have been extensively investigated using mathematical models. While smaller regulatory networks are modelled using continuous ODE based frameworks [41, 26], larger networks are modelled using discrete/logical approaches due to lack of scalability of ODE approaches in scenarios where network parameters may not be easily obtained [32]. While either mechanism may reveal atleast the most frequent steady state, a comparative analysis for larger networks, becomes computationally quite expensive. Hence, we offer an alternative modelling framework, which is continuous in nature yet parameter independent. This framework is computationally more efficient, offers more detailed analysis compared to the Boolean formalism and can reveal the underlying design principles of a network topology even for larger networks.

The scalability of our results can also be seen by considering the 22 and 26 node EMP networks that were simulated via RACIPE, wherein we were able to identify edge deletions/ nature changes that significantly reduce plasticity. Our previous work [28] has indicated that the reducing the number of positive feedback loops can reduce the plasticity associated with a network, but also showed that although some perturbations decreased the number of positive feedback loops to a comparable extent, but the corresponding fold change in plasticity was significantly different.

We hypothesized that this difference was due to them not taking into account the effect of negative feedback loops in the system, and were able to obtain a much stronger correlation using a weighted combination of negative and positive feedback loops in the network. As Boolean approaches fail to characterise the plasticity of large scale networks effectively, and ODE based methods such as RACIPE are too computationally expensive for large networks, such insights into the topological hallmarks provide a valuable tool to understand which links must be disrupted to curb plasticity, which could potentially be valuable in therapeutic applications, such as preventing metastasis by managing EMP.

We observed that EMP networks were relatively more dynamically robust as compared to the random networks we had generated of the same size. These results reinforce previous observations seen in other regulatory networks. The biochemical network involved in the establishment of segment polarity in Drosophila melanogaster has been shown to be robust against changes in initial values and rate constants of molecular interactions, enabling stable pattern formation [42, 43]. Similarly in a rewiring experiment for GRN of E. Coli, wherein new regulatory interactions were added to GRNs, [44] 95 % of these modified gene interaction networks were tolerated by the bacteria and some conferred selective advantages in particular environments. Our analysis of the structural robustness of EMP networks reveals similar trends, in terms of both plasticity and phenotypic distribution.

On further analysis of the topological properties that lend these networks dynamical and structural robustness, we found that robustness was associated with a larger number of positive feedback loops, and a smaller number of negative feedback loops. Moreover, the relative importance of each feedback loop was found to vary with both length and sign, across the different robustness measures. Multiple studies have modelled the evolution of biological networks, selecting for fitness [45], randomly generating GRNs using Markov Chain Monte Carlo [46] and using preferential attachment [33], finding that robust networks are rich in PFLs, and have relatively lower number of NFLs. Furthermore, the robustness offered due to these motifs can be evolutionarily stable [47] to edge perturbations, thereby possibly explaining the robustness of EMP in multiple biological contexts.

Our methods only investigate robustness on the gene regulatory level, however it is important to understand how the interactions between genes, proteins and metabolites contribute to robustness in a multi-layer heterogenous biological network [11]. Moreover, merely counting the number of feedback loops in the network is insufficient to capture the overall network topology, as these feedback loops can be coupled to varying degrees. Consequently, more sophisticated measures need to be developed that take into account the location and degree of interaction between different feedback loops in order to more effectively understand which loops must be disrupted for maximum effect. Other topological factors such as high interconnectivity, redundancy, degree distribution can also influence robustness [48, 49]. Despite these limitations, our results provide an integrated platform to understand the structural and dynamic robustness of EMP. Understanding the interplay between negative and positive feedback loops that lend these networks robustness, as well the mutations that can expose the fragility of these networks is crucial for drug design from a therapeutic perspective.

## Methods

### Random Network Generation

Random networks were generated uniformly across the set of all connected networks. We generated 100 random networks for each size, defined by the number of nodes in the network *n* = 4 – 10. The total number of edges in the network is decided before generation (*m*) with the constraint *m* ≥ *n* – 1, and picked uniformly from the binomial distribution:

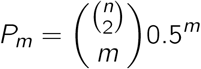

First a random spanning tree (a graph on the *n* nodes without any cycle with *n* – 1 edges) is generated by picking edges uniformly at random, and adding them if they don’t cause a cycle, ensuring that each node is connected with another node in the network. The remaining edges (*m* –*n* + 1) are then added randomly until the desired total number of edges is reached. The random networks generated for calculating perturbation JSD have an additional condition: each node of the network must have an outgoing edge. This is to ensure that there are no output nodes that unduly change the JSD despite the core network dynamics not changing.

The network sizes we analyze for each criteria of robustness are as follows

Avg. perturbation JSD: 4 –10

Avg. Fold change in plasticity (dynamic): 4 –8

Avg. Parameter variation JSD: 4 –8

Avg. Fold change in plasticity (structural): 4 –5

JSD between RACIPE and Cont. : 4 –8

Analysis for larger networks could be performed for criteria that only use the CSB, but those that also use RACIPE faced computational limitations, especially structural robustness in plasticity, as multiple perturbed networks need to be simulated for each network topology. The above datasets were generated in triplicates, and analysis performed in parallel. The corresponding measures are available in **Table S2**.

### Random Circuit Perturbation (RACIPE)

RACIPE [30] is a tool used to simulate gene regulatory networks (GRNs) in a continuous fashion. For a given GRN, RACIPE constructs a set of ODEs representing the interactions in the network. For a node i, let *A_i_*, and *i_i_*, denote the set of all activating and inhibiting input nodes to i respectively. Denote the expression of node i by *E_i_*,. The dynamics of node i is given by the ODE

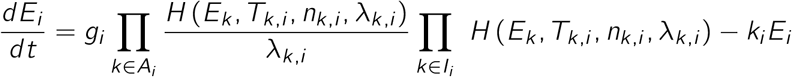

Where for node i, *T_k,i_* is the threshold value of the concentration *E_i_*, *g_i_*, is the production rate, *k_i_*, is the degradation rate, *n_k,i_*. is the hill coefficient, λ_*k,i*_, is the fold change. *H*^+^ is called the shifted hill function, given by:

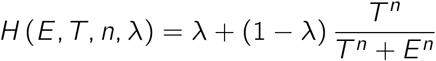

RACIPE simulates these ODEs by randomly sampling parameter sets from a pre-determined range of parameters uniformly. These parameter ranges were estimated from BioNumbers [50]. For each parameter set, the ODEs are simulated for multiple initial conditions. The parameters used for each such set of simulations and the corresponding stable states obtained in that domain are the outputs obtained.

### Discretising RACIPE data, calculating stable state frequencies and stability analysis

For each parameter set sampled by RACIPE, the different initial conditions generated converge to a number of steady states. Each steady state obtained from RACIPE is assigned a weight, equal to the fraction of initial conditions that converge to the steady state for the corresponding parameter set, resulting in *M* × (*N* + 2) table, where *N* is the number of nodes in the network and *M* is the number of steady state-parameter set combinations. Since each parameter can have more than 1 steady states, *M* >= *N*. The first column showing the parameter ID, next *N* columns showing the expression level of each node of the network and the last column showing the weight for the corresponding steady state. The node expression values are converted to weighted z-scores by scaling them about their means:

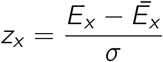

where the steady state expression vector of a node x across all steady state-parameter combinations is given by by *E_x_*, *Ē_x_* is the weighted mean of the steady state expression of node *x*, and σ is the weighted standard deviation of the expression level. The weighted z-scores are then binarised by assigning a value of 1 for positive and 0 for negative weighted z-scores respectively. Hence, each steady state is a string of 1s and 0s, with the number of nodes as the length. While calculating the frequency of a stable state for a parameter set, we weight them by the fraction of initial conditions converging to that state.

### Boolean Model

The Boolean algorithm devised by Font-Clos et al [32] is used here. Nodes have binary values of −1(low), 1(high). If *x_i_*, is the state of a node, the activation *A_i_*, is given by

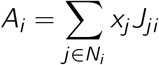

where *N_i_*, is the set of nodes activating/inhibiting node *i*, and *J_ji_* is the sign of the interaction. Nodes are updated asynchronously by choosing a node uniformly at random at every time step, and changing it to 1 if the node’s activation is positive, and −1 if it is negative. If the activation is 0, then we don’t update the node.The final steady states are represented in a 0(low), 1(high) format, for ease of readability.

### Continuous State-Space Boolean (CSB) Formalism

The CSB formalism is based on the Boolean model by Font-Clos et al [32], but with a continuous state space. In this model, the expression values of the nodes in a GRN are continuous real values restricted between [−1, 1] (1 corresponding to ON and −1 corresponding to OFF). The update rule is formulated to be similar to the euler integration method. The rate of change in the expression level is modelled as a sigmoidal function, as follows:

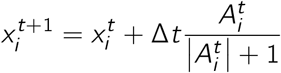

Where 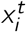 is the expression of node *i* at time *t*, 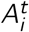 is the activation of node *i* at time *t*, defined as:

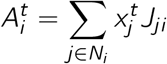

*N_i_*, is the set of nodes activating/inhibiting node *i*, *J_ij_* is the sign of the edge from node *j* to node *i*. Δt = 0.1 corresponds to a small discrete timestep. Upper and lower bounds of −1 and 1 are enforced on the expression levels during the simulations, such that if 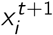 goes below −1 (or goes above 1), 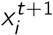 is set to −1 (or 1). A state at time *t* is classified as stable or unstable based on the following condition:

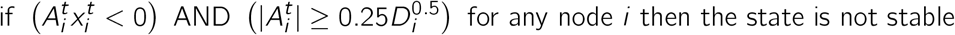

where *D_i_* is the indegree of node *i*. The condition can be described as follows: if the activation of any node has an opposite sign as the expression level of the node (thereby pushing the node towards the opposite sign), and if the activation is large enough for the push to be of any consequence, then the state is unstable. The lower bound for the activation threshold 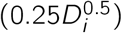 is of the order of the standard deviation of activation of a node, if input nodes were randomly initialised. Essentially, we only consider the state to be unstable if the sign of some node doesn’t match its activation, AND the activation is sufficiently large in modulus. The networks are simulated until a stable state is reached. The stable state values are then binarised as 1 for positive stable state, and 0 otherwise.

### Fold change in plasticity

Fold change in plasticity is used to measure the degree of change in plasticity of the wild type upon a perturbation, either dynamic or structural. If the plasticity of the wild type network is *p*_1_ and the plasticity of the perturbed network is *p*_2_, then:

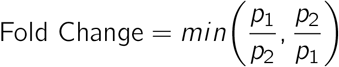

Consequently, the fold change ranges from 0 to 1, and a higher value indicates that the perturbed network has plasticity closer to the wild type. Hence, looking at the average fold change due to perturbation (structural, or kinetic parameter variation) gives a good estimate of the robustness in plasticity of the network.

### Jensen Shannon Divergence

The difference between two phenotypic distributions can be quantified by the JSD metric. For two discrete frequency distributions P(x) and Q(x), we define:

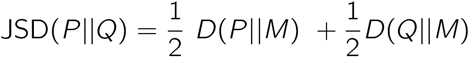

Where 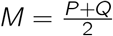, and D is the Kullback-Leibler divergence, given by:

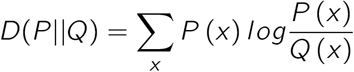

JSD varies from 0 to 1 when base to is used to calculate the logarithm in D, with 0 indicating that the distributions are identical, and 1 indicating that the distributions have no overlap. The JSD values were calculated using the jensenshannon function in the scipy.spatial.distance module (Python 3.8).

### Comparison of parameter variation curves for plasticity

In order to understand how important of a role the hill coefficient plays in the kinetic variation of a network, we expanded/shrunk the parameter space in 3 different regimes: All parameters varied simultaneously, only the hill coefficient being varied, everything but the hill coefficient being varied. The absolute area between the control (all parameters) and the 2nd and 3rd regimes were then computed in order to quantify the closeness of the parameter variation schemes, with a lower value indicating that the curves are more similar.

### Percent error in Phenotypic distribution

In order to estimate the degree of variance in distribution across multiple simulations, the following metric was employed. For a given network *N*, suppose it has stable state frequencies *S*(*i*). The simulation is then done with 10 different runs, from which the error in *S*(*i*), *E*(*i*) is found for each state by finding the standard deviation. The percent error for that simulation is then defined to be

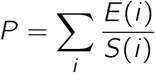

This is repeated in triplicates, and the average value of *P* is reported as the fraction error.

### Calculating the number of Feedback Loops

The number of feedback loops in a given network were calculated using the NetworkX package[51]. We first count the number of cycles in the network using the *simple_cycles* function, then traverse through each cycle multiplying the edge signs up (1 for activation and −1 for inhibition). The cycle is classified as a positive feedback loop if the product is positive, and negative otherwise.

### Weighting of Feedback Loops

There are 2 types of weighting that we use: weighting by length, and weighting according to sign. First, each feedback loop in the network is weighted by the inverse of its length, to obtain the total number of weighted (by length) positive feedback loops (WPFL) in the network.

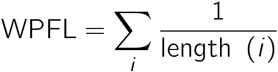

Where the sum is over all positive feedback loops. The number of weighted negative feedback loops (WNFL) is calculated similarly. We then assign differential weights to feedback loops on the basis of their sign. Because negative feedback loops were generally found to be negatively correlated with our robustness metrics, they receive a negative weight, normalised to −1, and PFLs receive a positive weight (a). In other words, the total number of weighted feedback loops in the system (WFL) is given by

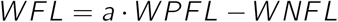

Similarly, we can weight the feedback loops just by their sign (not length) to obtain SWFL, the number of signed weighted feedback loops.

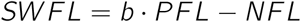

### Fraction of positive cycles (unweighted and weighted)

The fraction of positive cycles (FPC) in the network is simply given by

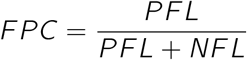

Similar to the above, one can also weight the cycles (by length and sign) when calculating the fraction of weighted positive cycles, (FWPC)

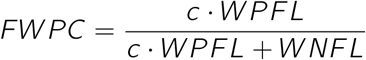

So, as the WNFLs increase, the fraction decreases, thereby maintaining negative correlation between negative feedback loops and robustness as in the previous case.

### Optimal weights for a given indicator of robustness

For a given indicator of robustness (e.g., average Perturbation JSD), the optimal weight *a* in the above 2 metrics (number of WFLs or FWPC, the *x* variable) for the perturbed networks is found, that maximises the correlation with the robustness indicator (the *y* variable). This is done using the curve_fit module in SciPY package in Python 3.8.

### Simulation Parameters

Unless mentioned otherwise, the number of simulations used to estimate phenotypic distributions or plasticity in Boolean/RACIPE/Continuous formalisms is chosen to be 10000 times the number of nodes in the network. For RACIPE, the number of initial conditions for each parameter set was 100. The effect of initial conditions on the population level measures such as plasticity and phenotypic distribution is minimal, as shown in our previous study [28]. Error bars are obtained by performing the simulations in triplicates. RACIPE was run with the default parameters, with only the number of parameter sets being varied across networks. The topofiles for each network analyzed as well as .ids files containing the order of nodes in the state are present in the Github repo.

### Statistical tests and functions

All correlation analysis was done using Spearman correlation method using the scipy.stats.pearsonr module. The corresponding statistical significance values are represented by ‘*’s, to be translated as: * : *p* < 0.05, ** : *p* < 0.01, *** : *p* < 0.001. Unpaired T-test for the violin plots was performed using the scipy.stats.ttest_ind function, with significance being reported as above. Regression bands (95% CI) for correlation plots were plotted using seaborn.regplot.

## Data and Code Availability

The raw data generated for this study, is available at https://bit.ly/3iHZBSd. Derived datasets and code supporting the current findings, including the codes for continuous boolean model, and codes to generate SBML models of the networks are available on the github page: https://github.com/csbBSSE/Robustness_project

## Acknowledgements

AH and AM are supported by the KVPY fellowship awarded by Department of Science and Technology (DST), Government of India. KH is supported the the Prime Ministers Research Fellowship (PMRF). MKJ is supported by the Ramanujan Fellowship (SB/S2/RJN 049/2018) awarded by the Science and Engineering Research Board (SERB), DST, Government of India. Mr. Atchuta Srinivas Duddu is acknowledged for artwork **(Fig 1a,e, Fig 3a)**.

## Author Contributions

MKJ conceived and supervised the research. AH, AM and KH performed the research and analyzed the results. All authors prepared the manuscript.

## Additional Information

Correspondence and requests for materials should be addressed to MKJ.

## Supplementary

**Supplementary Figures**

**Fig. S1.**
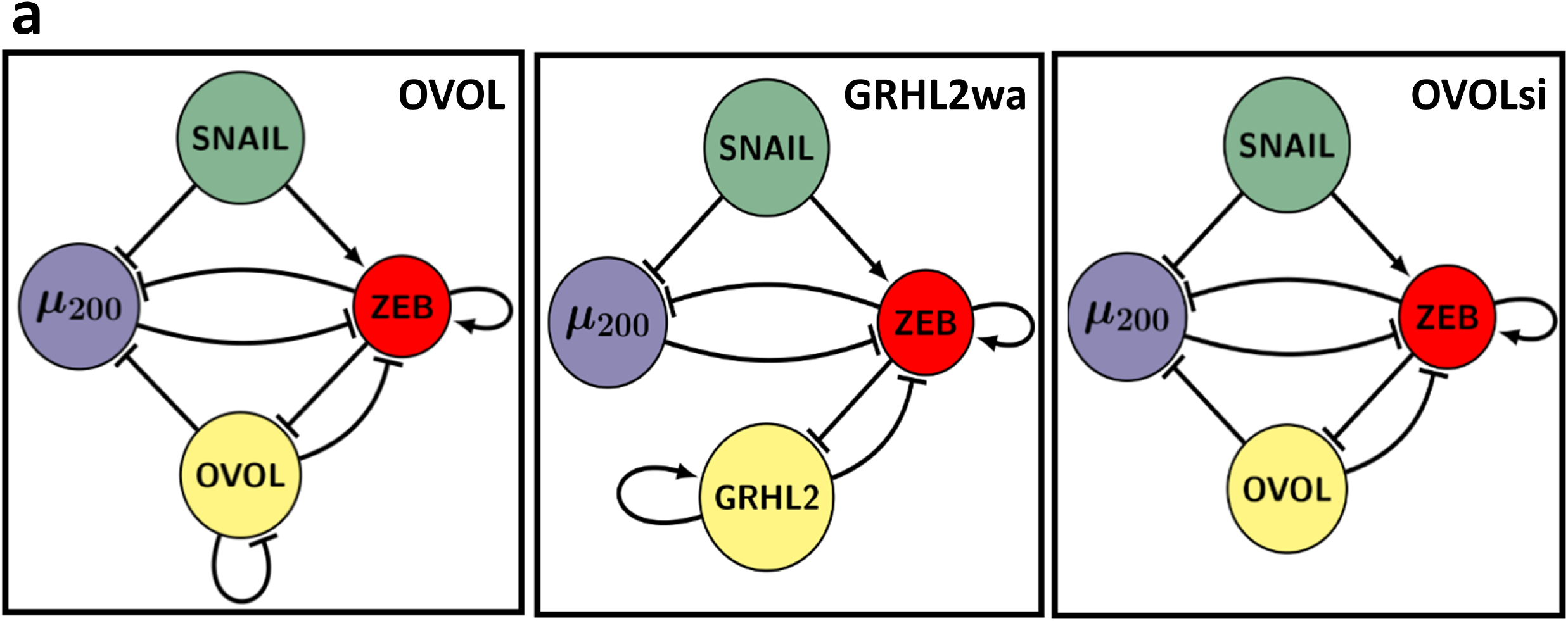
**(a)** From left to right: OVOL (4 nodes, 9 edges), GRHL2wa (4 nodes, 8 edges), OVOLsi (4 nodes, 8 edges).

**Fig. S2.**
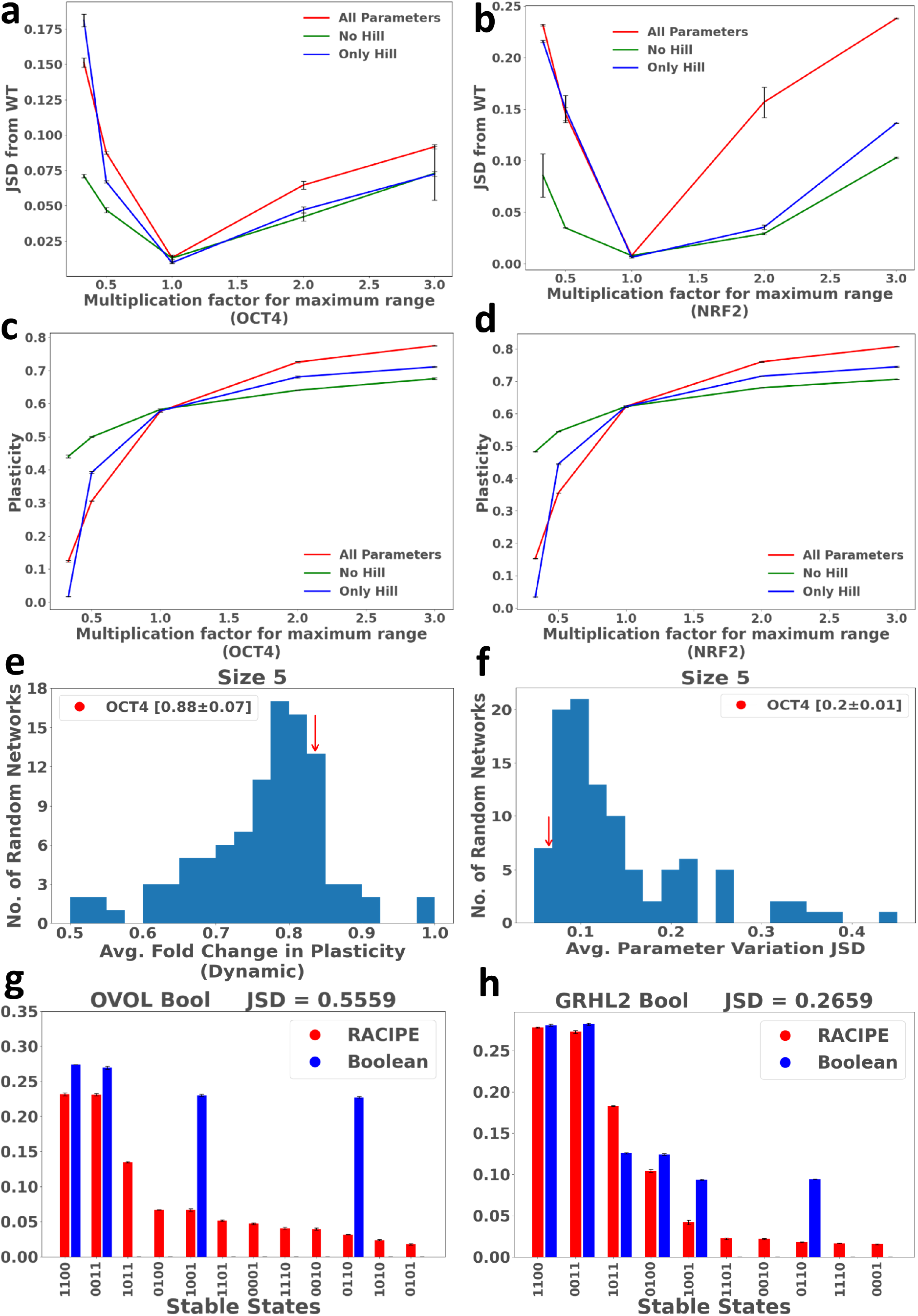
**(a,b)** Parameter space JSD variation plots for OCT4 and NRF2. **(c,d)** Similar to **a,b**, but for plasticity. **(e,f)** Distribution of average parameter variation JSD and fold change in plasticity for networks of size 5, with OCT4 marked. **(g,h)** Phenotypic distributions for OVOL and NRF2 networks, obtained using RACIPE and Boolean formalisms, compared using JSD. Error bars for phenotypic distributions found using the STD of 3 triplicates.

**Fig. S3.**
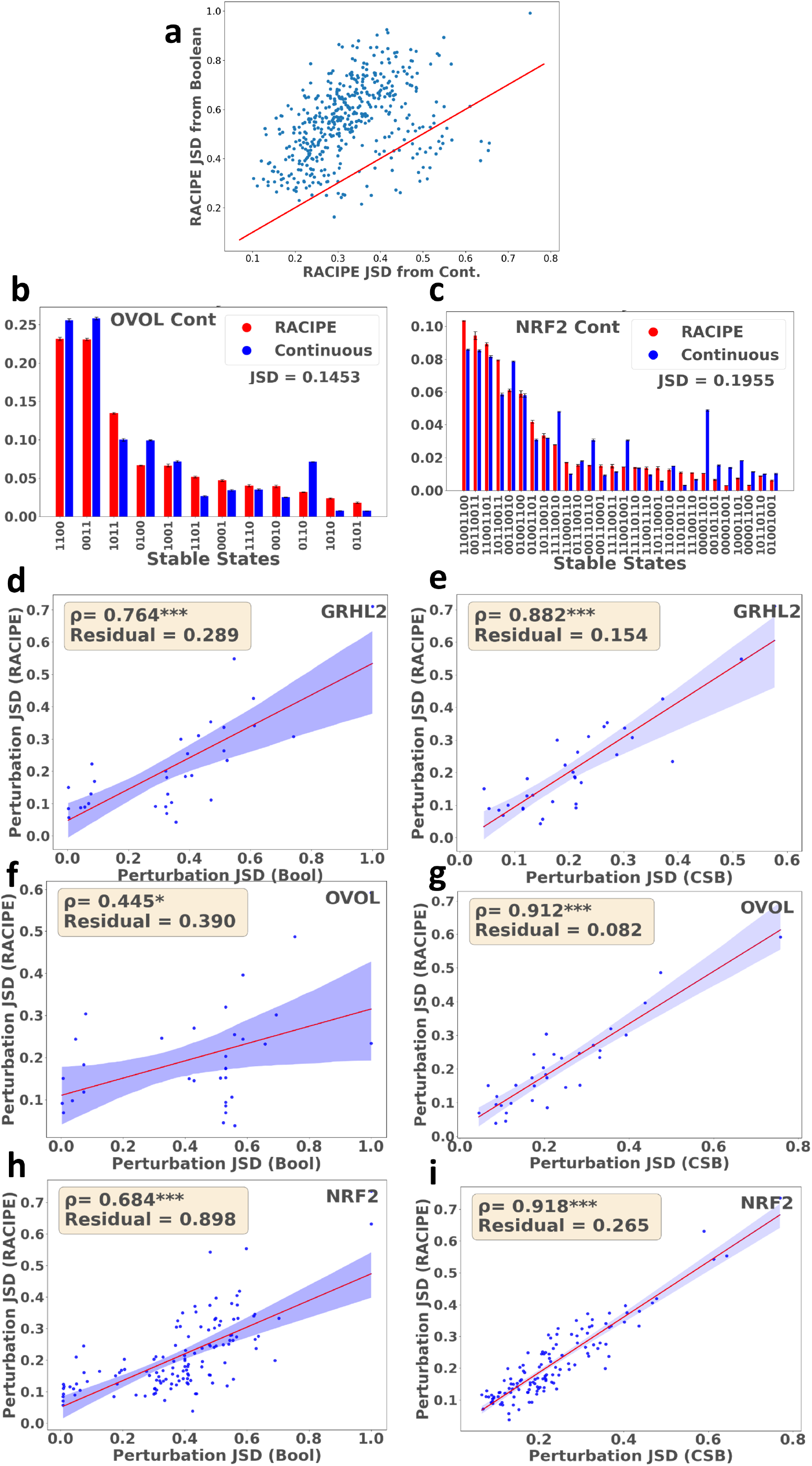
Comparison of JSD of the phenotypic distributions obtained using RACIPE from that obtained using Boolean formalism (y axis) and Continuous formalism (x axis). The red line is the x=y line. **(b,c)** Comparison of the phenotypic distributions obtained from RACIPE and Continuous model for OVOL and NRF2, using JSD. **(d,e,f,g,h,i)** Comparison of Boolean and Continuous model for the correlation in perturbation JSD with that of RACIPE for GRHL2, OVOL and NRF2 networks. Error bars for phenotypic distributions found using the STD of 3 triplicates.

**Fig. S4.**
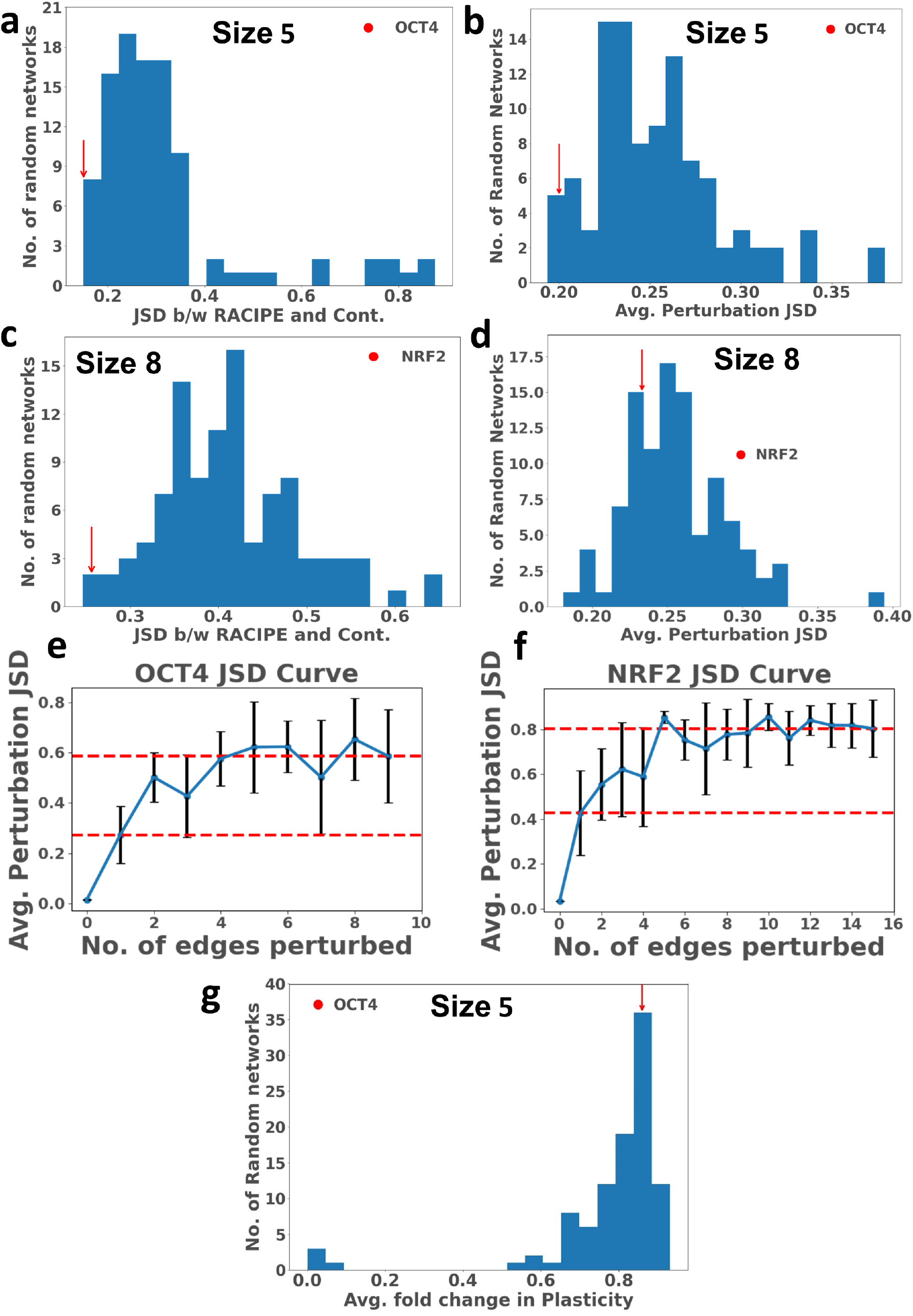
**(a)** Distribution of the JSD between RACIPE and Continuous model phenotypic distributions (dynamic robustness measure) for random networks of size 5, with OCT4 marked. **(b)** Same as **a**, but for average perturbation JSD. **(c,d)** Same as **a,b**, but for networks of size 8. **(e,f)** Multi edge perturbation JSD plots for OCT4 and NRF2. 10 random edge perturbations (of corresponding size) are performed for each size, with error bars being the STD. Horizontal red lines mark the average perturbation JSD due to a single edge perturbation, as well as E many perturbations. **(f)** Same as **a**, but for average fold change in plasticity.

**Fig. S5.**
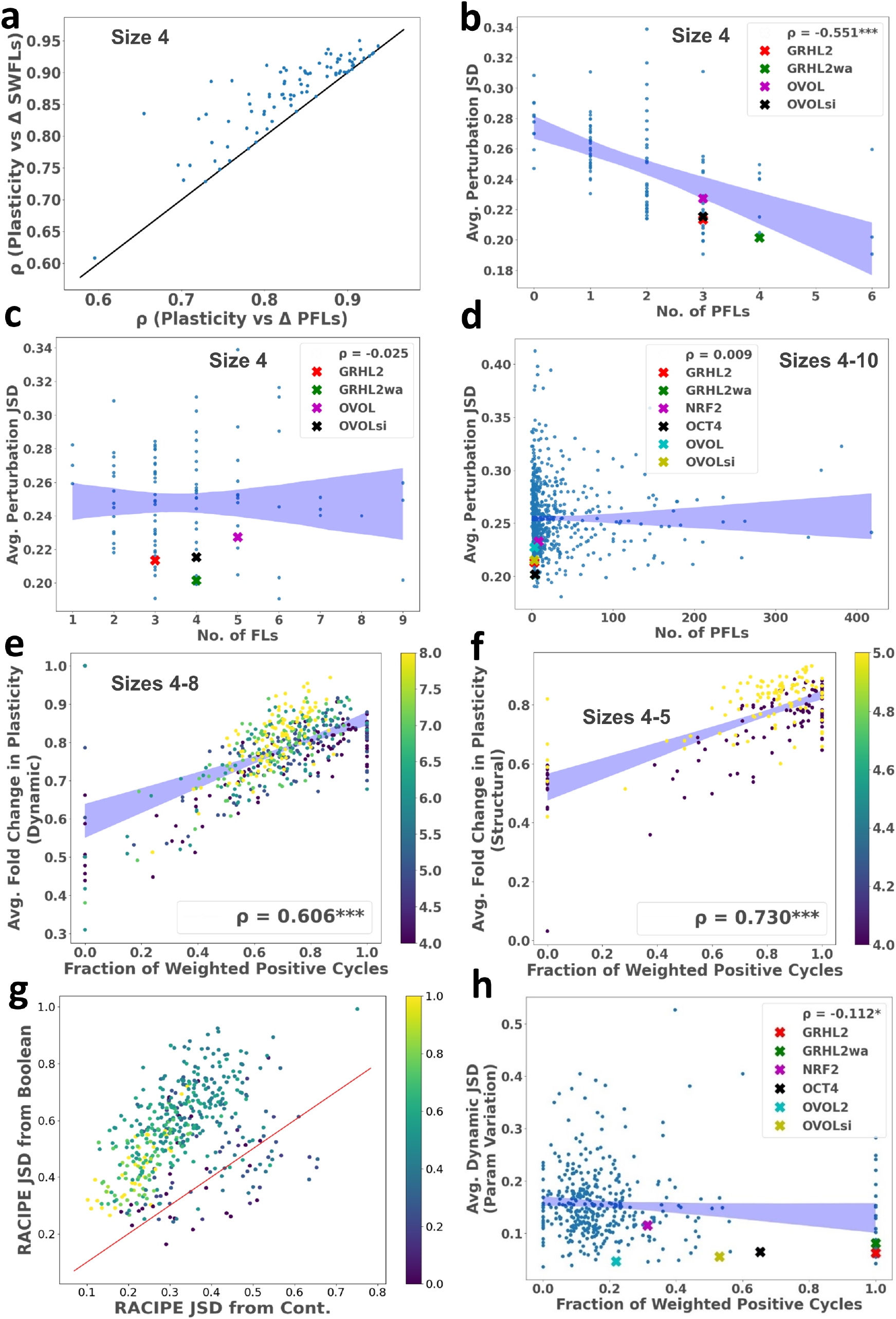
**(a)** The correlation between fold change in plasticity and cycle sum (PFLs - NFLs) and number of weighted feedback loops (by sign) was found for each network of size 4 and plotted as a scatter plot, along with the *x* = *y* line. **(b)** Scatter plot of the average perturbation JSDs from WT vs No. of positive feedback loops for networks of size 4, with the corresponding WT networks marked. **(c)** Same as **b**, but for networks of sizes 4-10 combined. **(d)** Dynamic fold change in plasticity vs fraction of weighted positive cycles for networks of size 4-8, coloured by size. **(e)** Average fold change in plasticity vs fraction of weighted positive cycles for networks of size 4-5, coloured by size **(f)** Avg. parameter variation JSD vs fraction of weighted positive cycles for networks of size 4-8, with the corresponding WT networks marked. **(g)** Comparison of JSD of the phenotypic distributions obtained using RACIPE from that obtained using Boolean formalism (y axis) and Continuous formalism (x axis). The red line is the *x* = *y* line. The networks are coloured by the fraction of positive cycles (unweighted).

**Fig. S6.**
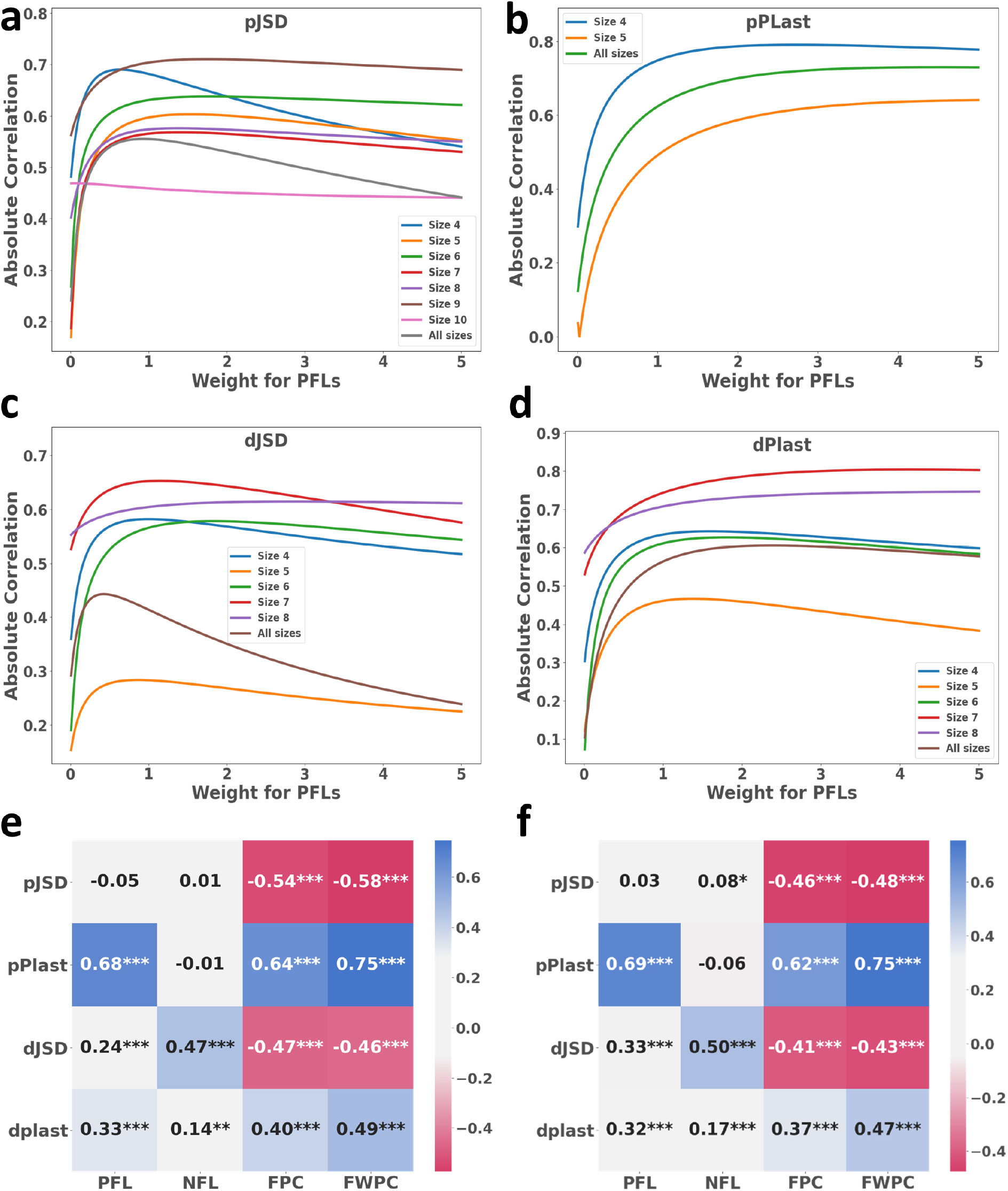
**(a)** Line plot indicating the correlation with average pJSD with varying weight assigned to PFLs in the FWPC metric, for different sizes. **(b,c,d)** Same as **a**, but for pPlast, dJSD, and dPlast respectively. **(e,f)** Heatmap depicting the correlation between robustness measures and network metrics, for 2 sets of 100 random networks each. Notice the similarity between the correlation values between the two heatmaps and that in **Fig 5i**

**Fig. S7.**
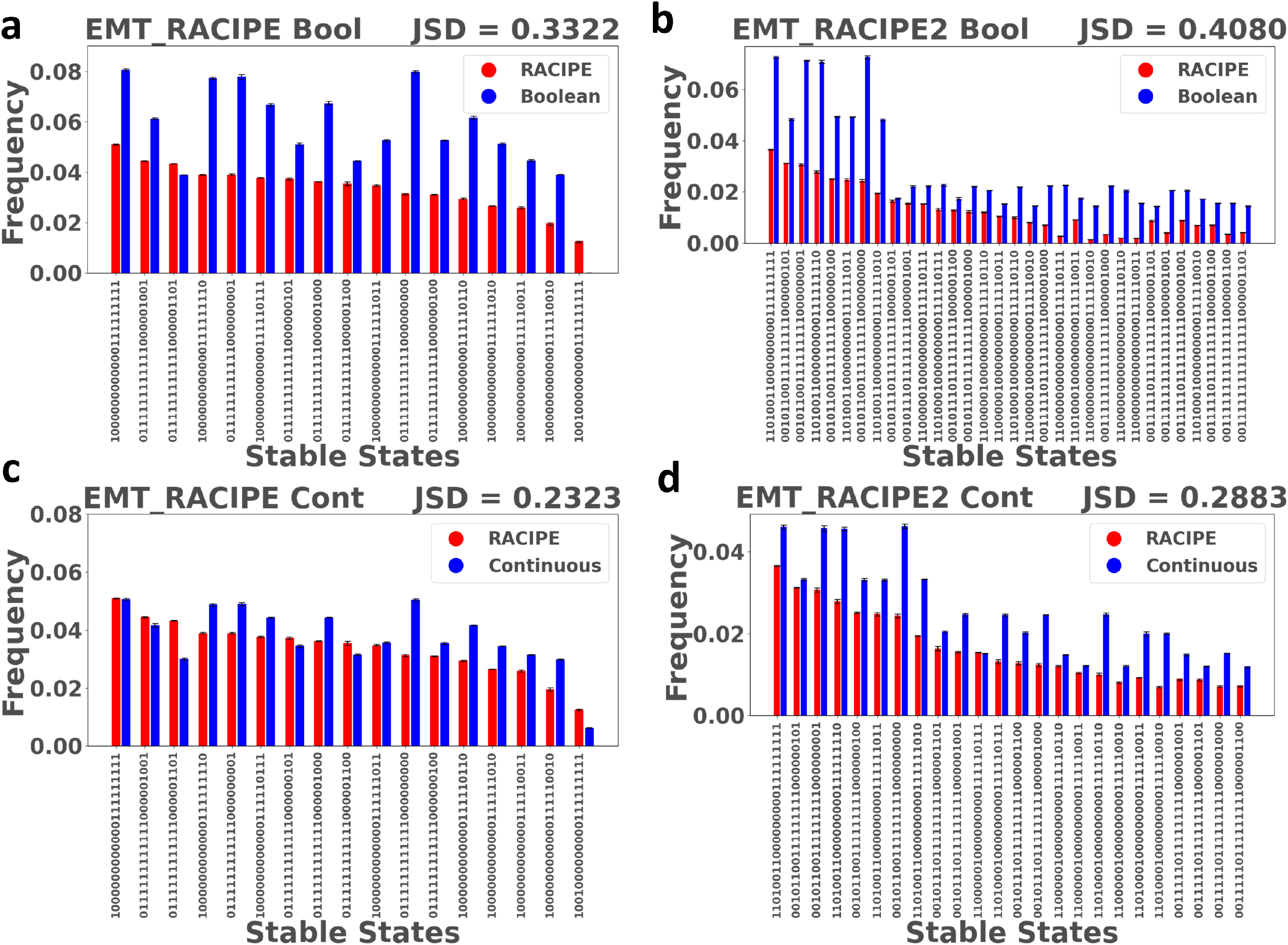
**(a,b)** Boolean vs Racipe phenotypic distributions for EMT_RACIPE and EMT_RACIPE2, compared using JSD. **(c,d)** CSB (Continuous State Space Boolean) vs Racipe phenotypic distributions for EMT_RACIPE and EMT_RACIPE2, compared using JSD. Error bars for phenotypic distributions found using the STD of 3 triplicates.

